# Lipid Metabolism Remodeling in Human Cardiomyocyte Differentiation and Maturation

**DOI:** 10.1101/2025.10.21.682321

**Authors:** Haite Tang, Raina Liu, Zhiqiang Xiong, Qiqi He, Gang Lu, Wai-Yee Chan, Wuming Wang

## Abstract

Cardiac development involves extensive metabolic remodeling, yet dynamic changes of endogenous lipid species during human cardiomyocyte (CM) differentiation remain poorly characterized. In this study, we systematically performed high-coverage lipidomic profiling across five consecutive CM differentiation stages, using high-performance liquid chromatography. *MYL2* expression confirmed ventricular lineage specification in late-stage CMs. Ceramide levels peaked at the cardiac progenitor cell (CPC) stage, suggesting a stage-specific association with the mesendoderm (MES)-to-CPC transition. Meanwhile, lipid categories such as monosialodihexosylganglioside, phosphatidylcholine (PC), lysophosphatidylcholine, and lysophosphatidic acid were markedly elevated in mature CMs. Integrated transcriptomic and chromatin accessibility analyses further delineated the temporal regulatory landscape underlying stage-dependent lipid remodeling during cardiogenic differentiation. Functional assays revealed that PC promotes the expression of mature CM marker genes and improves the CM viability. Collectively, these findings establish a comprehensive, high-resolution dynamic lipidomic atlas of *in vitro* hESC-derived CM differentiation, providing a valuable resource for understanding lipid-associated mechanism in heart development.

## INTRODUCTION

Cardiovascular diseases remain the foremost cause of global mortality, accounting for 31% of all deaths worldwide^1^. Given the heart’s limited regenerative capacity, conventional therapies are insufficient to replace lost cardiomyocytes (CMs), underscoring the urgent need for innovative regenerative strategies. Human pluripotent stem cells (PSCs), with their ability to differentiate into CMs, offer a promising resource for cardiac regeneration^2^. Transcriptomic analyses have revealed enrichment of lipid-associated factors (e.g., *LIPTER*) in iPSC-derived CMs and aberrant regulation of lipid metabolism pathways in peripartum cardiomyopathy-specific iPSC-CMs versus healthy controls^3,4^. These findings indicate that transcriptomic profiling of iPSC-CMs effectively identifies functional lipids driving cardiogenesis. While the epigenome and proteome of mesodermal cells, cardiac mesoderm, immature CMs, and mature CMs derived from PSCs have been characterized to identify key regulators of CM differentiation^5,6^, metabolomic insights—particularly lipidomic profiling—remain limited, especially when integrated with transcriptomic data.

Lipid metabolism remodeling is crucial in heart development, supporting structural maturation and metabolic transitions essential for function. Lipid-derived signaling molecules such as sphingosine 1-phosphate (S1P) and lysophospholipids regulate CM proliferation, differentiation, and survival^7,8^. S1P and lysophosphatidic acid (LPA) enhance CM generation when administered during the early stage of differentiation^9^. Lipoprotein lipase deletion impairs embryonic heart development by inhibiting CM proliferation and disrupting ventricular chamber formation^10^. Altered lipid droplet metabolism is linked with congenital heart defects and cardiac arrhythmias^11–13^.

Fatty acid (FA) promotes a metabolic switch toward mitochondrial oxidative phosphorylation, accompanied by reduced dependence on glycolysis and glutamine metabolism during CM maturation^14,15^. Notably, FA oxidation inhibition enhances heart regeneration in adult animal models^16^, and sphingolipid (SL) metabolism plays a critical role in CM proliferation and regeneration^8^.

Metabolomics has identified metabolic markers indicative of maturation in human PSC- derived CMs^17^. During early cardiac development, CMs rely predominantly on glycolysis; with ventricular thickening and increased energy demand, FA oxidation becomes the main ATP source, contributing 70–90% at rest^18–20^. CMs acquire FAs via circulating lipoprotein uptake and store them as triglycerides^21^. FA supplementation promotes maturation of PSC- derived CMs^22^.

This rapidly evolving field highlights the intricate interplay among lipid metabolism, cardiac development, and disease. In this study, we performed integrative multi-omics analyses across five developmental stages to delineate stage-specific lipidomic remodeling. We further validated the link between phospholipid metabolism and CM developmental states using functional supplementation assays. Collectively, our findings provide a comprehensive framework for understanding metabolic transitions during in vitro differentiation and maturation of hESC-derived CMs, with potential implications for future translational applications.

## RESULTS

### Temporal Lipidomic Profiling Reveals Stage-Specific Signatures During Human Embryonic Stem Cell (hESC)-to-Cardiomyocyte (CM) Differentiation

To investigate lipid remodeling during early human heart development, we induced the differentiation of human ESCs into CMs via small-molecule WNT modulation, recapitulating key stages of fetal cardiac development. Five key stages were selected for lipidomic and transcriptomic analyses: ESC (Day 0), mesoderm (MES, Day 3), cardiac progenitor cells (CPCs, Day 6), immature CMs (Day 10), and mature CMs (Day 30) (**Figure** 1A, B). Ventricular myosin regulatory light chain (MLC2v), encoded by *MYL2*, is a classical marker of ventricular lineage CMs and a hallmark of maturation. iPSC-derived MLC2v-positive CMs have been recognized as mature cells^23,24^, and abundant MLC2v expression has been reported in iPSC-derived CMs by day 20–30 post-differentiation^23–25^. In this study, RNA-seq analysis and immunofluorescence (IF) staining confirmed significant MLC2v enrichment in day-30 CMs (**Figure** S1A, B). Additionally, using an iPSC clonal line with endogenous *MYL2*-mEGFP tagging, we observed robust MLC2v-EGFP signals in spontaneously beating day-30 CMs (**Video** S1). Quantitative analysis of low-magnification IF fields confirmed that cTNT-positive CMs accounted for 90%-94% of the total cell population at Day-10 CMs, while MLC2v-positive ventricular-like CMs accounted for 88%-95% of the total cell population at Day-30 CMs, indicating progressive ventricular lineage specification and high CMs purity throughout late differentiation stages (**Figure** S1C, D).

**Figure 1.**
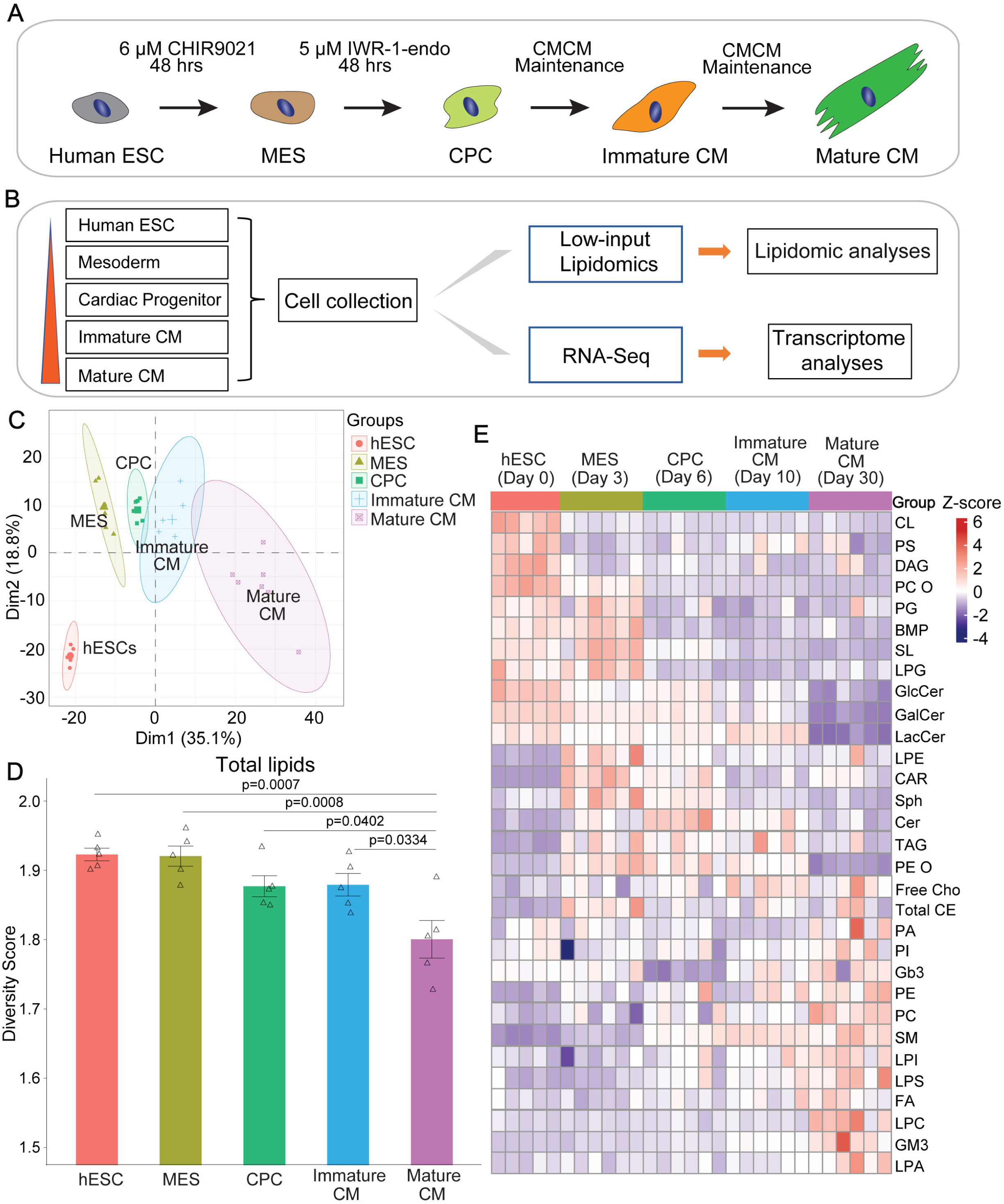
Lipidomics signatures remodeling during hESC-derived CM differentiation (A) Schematic overview of the experimental workflow for directed differentiation of hESCs into CMs via the sequential small-molecule modulation of WNT signaling. CMCM, CM Culture Medium. (B) Detailed schematic workflow illustrating stepwise sample collection at five key developmental time points, followed by systematic lipidomic quantification and transcriptomic RNA sequencing integration. (C) Principal component analysis (PCA) score plot based on global lipidomic profiling data, demonstrating clear separation and tight intra-group clustering across five consecutive differentiation stages: ESCs (Day 0; *n* = 9), mesoderm (MES, Day 3; *n* = 9), cardiac progenitor cells (CPCs, Day 6; *n* = 10), immature CMs (Day 10; *n* = 7), and mature CMs (Day 30; *n* = 7). (D) Quantitative evaluation of global lipid molecular diversity across the entire CM differentiation time course. Lipid diversity was calculated using the Shannon-Wiener entropy index: Shannon index = −∑Pi × log(Pi), where Pi represents the relative molar fraction probability of each individual lipid species within the total lipid pool. (E) A global heatmap illustrates the normalized relative molar abundances of all major annotated lipid categories across five staged CM differentiation samples. Color scale represents global z-scores of relative molar fractions across all detected lipid species. All lipid profiles were derived from the same biological replicate cohort as presented in (C). Abbreviations: CL, cardiolipins; PS, phosphatidylserine; DAG, diacylglycerol; PC O, Alkyl phosphatidylcholine; PG, phosphatidylglycerol; BMP, bis(monoacylglycero)phosphate; SL, sphingolipid; LPG, lysophosphatidylglycerol; GlcCer, glucosylceramide; GalCer, galactosylceramide; LacCer, lactosylceramide; LPE, lysophosphatidylethanolamine; AC(CAR), acylcarnitine; Sph, sphingosine; Cer, ceramides; TAG, triacylglycerol; PE O, Alkyl phosphatidylethanolamine; Free Cho, free cholesterol; Total CE, total cholesteryl ester; PA, phosphatidic acid; PI, phosphatidylinositols; Gb3, globotriaosylceramide; PE, phosphatidylethanolamine; PC, phosphatidylcholine; SM, sphingomyelin; LPI, lysophosphatidylinositol; LPS, lysophosphatidylserine; FA, free fatty acid; LPC, lysophosphatidylcholine; GM3, monosialodihexosylganglioside; LPA, lysophosphatidic acid. All quantitative data are presented as mean ± standard error of the mean (SEM) from independent biological replicates (*n* = 5–6). Statistical differences in pairwise diversity comparisons between adjacent differentiation stages were evaluated using one-way ANOVA (^∗∗∗^*p* < 0.001, ^∗∗^*p* < 0.01, and ^∗^*p* < 0.05).

To further characterize the acquisition of maturation-associated structural features, we performed co-IF staining for ACTN2 (a sarcomeric Z-line marker) and TIM23 (a key component of the mitochondrial inner membrane). Confocal imaging revealed that Day-30 CMs developed highly organized, parallel-aligned myofibrils with clear periodic striations corresponding to alternating Z-lines, indicative of well-assembled sarcomeric architecture at a late stage of culture. In parallel, Day 30 CMs displayed abundant, elongated mitochondria, consistent with enhanced cellular maturation (**Figure** S1E). Electrophysiological analysis using the Maxwell Biosystems platform for Day-30 CMs revealed an extracellular spike pattern, indicating physiological maturity and electromechanical competence of Day-30 CMs (**Figure** S1F).

These data confirm reliable stage-specific differentiation and high CM purity from day 10 onward, supporting the use of this system for stage-resolved lipidomic profiling.

We applied a low-input trace lipidomics method adapted from published protocols^26^ to maximize lipid species detection (**Figure** 1B). Principal component analysis (PCA) demonstrated distinct clustering by differentiation stage, indicating high sample reproducibility and stability (**Figure** 1C). Shannon diversity index analysis revealed a gradual decrease in lipid diversity throughout differentiation (**Figure** 1D). Across samples, 31 major lipid classes with 781 subclasses were identified (**Figure** 1E).

Lipids were clustered based on dynamic trends during differentiation (**Figure** 1E, S2A-H). Cardiolipins (CLs), diacylglycerols (DAGs), phosphatidylcholine oleic acid subtypes (PC Os), phosphatidylglycerols (PGs), glucosylceramides (GlcCer), and galactosylceramides (GalCer) peaked at the ESC stage and declined thereafter (**Figure** S2A, B). Phosphatidylserines (PSs), phosphatidic acids (PAs), globotriaosylceramides (Gb3), and lactosylceramides (LacCer) exhibited an initial decline followed by an increase during differentiation (**Figure** S2C).

Sphingolipids (SLs), acylcarnitines (ACs), lysophosphatidylethanolamines (LPEs), sphingosines (Sphs), bis(monoacylglycero)phosphates (BMPs), total cholesteryl esters (CEs), and phosphatidylethanolamine oleic acid subtypes (PE Os) peaked at the MES stage (**Figure** S2D, E). Free cholesterol (Free Cho), triacylglycerols (TAGs), ceramides (Cers), and lysophosphatidylglycerols (LPGs) peaked at CPC or immature CM stages (**Figure** S2D, E). Fatty acids (FAs), PEs, phosphatidylinositols (PIs), sphingomyelins (SMs), monosialodihexosylgangliosides (GM3), PCs, lysophosphatidylcholines (LPCs), lysophosphatidylinositols (LPIs), lysophosphatidylserines (LPSs), and lysophosphatidic acids (LPAs) generally increased progressively during differentiation (**Figure** S2F-H).

### Dynamics of Monotonically Downregulated Lipid Subclasses During Early CM Differentiation

Among the major lipid classes identified, those predominantly functioning as cell membrane components include BMP, free cholesterol (Free Cho), GalCer, Gb3, GlcCer, LacCer, PC, PC O, PE, PE O, PG, SL, SM, CL, AC, PA, Cer, GM3, PS, and PI, with mitochondrial membranes notably enriched in PE and CL^27–29^. Lipid classes such as AC, PA, Cer, GM3, PS, PI, DAG, LPA, LPC, LPE, LPG, LPI, LPS, and Sph act primarily as signaling molecules^27,30–34^. FA, CE, and TAG mainly serve as energy storage molecules^35,36^. Additionally, LPA has been implicated in regulating MES differentiation^37^, whereas LPC influences cationic currents in guinea pig atrial cells^38^. CL biosynthesis is essential for postnatal cardiomyocyte maturation, as it supports mitochondrial cristae maturation and the accumulation of oxidative phosphorylation complexes^39^^51^. To profile the dynamic abundance changes of each lipid class during CM differentiation, we systematically conducted relative abundance of all lipid subclasses detected.

CL, a unique phospholipid with four fatty acid chains and strong hydrophobicity, primarily localizes to mitochondrial membranes, especially the inner membrane^40–42^. Both total CL relative abundance and most of the 54 identified CL species peaked at the human ESC stage, followed by a marked decline through to mature CMs (**Figure** 2A, S2A). Seventeen DAG subclasses, intermediates in lipid biosynthesis, showed maximal relative abundance at the ESC stage and gradually decreased during differentiation (**Figure** 2B). DAG 36:4 (18:2/18:2), DAG 36:3 (18:2/18:1), DAG 36:1 (18:1/18:0), and DAG38:4 (18:0/20:4) were relatively enriched in mature CMs, with DAG 38:4 (18:0/20:4) showing the most prominent increase (**Figure** 2B). BMP, a biosynthetic precursor of CL, displayed abundance patterns parallel to CL, suggesting a potential association with mesoderm-stage lipid remodeling (**Figure** 2C). PG species, direct precursors of CL, demonstrated stage-specific enrichment; four species - PG 36:3 (16:0_20:3), PG 36:4 (16:1_20:3), and PG 38:3 (18:0_20:3) - were enriched in early differentiation, whereas others peaked in mature CMs (**Figure** 2D). In short, PG species displayed stage-specific distribution, with highly unsaturated species more abundant at later differentiation stages.

**Figure 2.**
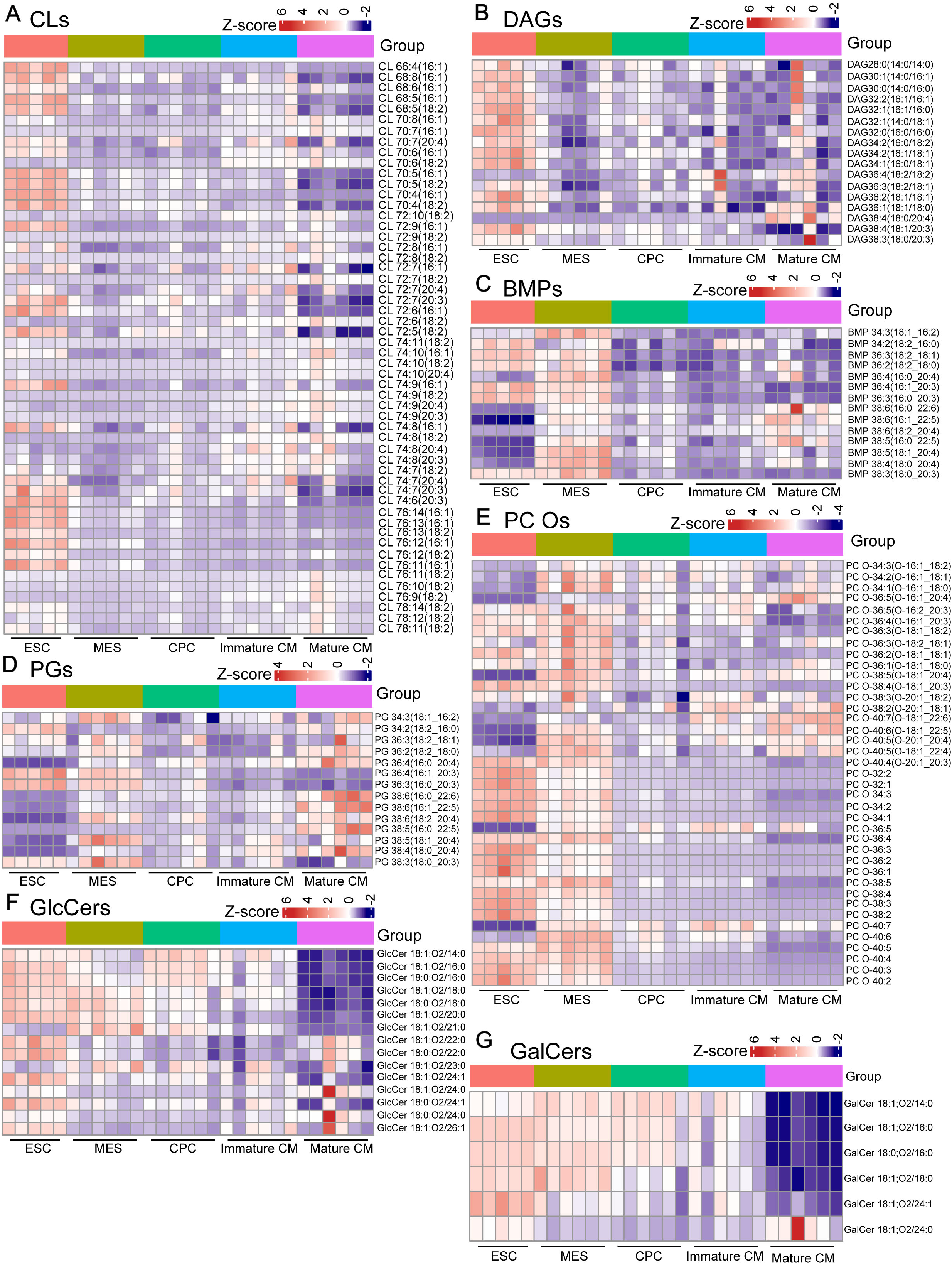
Dynamic alterations in monotonically downregulated lipid subclasses during hESC-derived CM differentiation (A) –(G) Stage-resolved heatmap overview illustrating temporal abundance dynamics of all lipid subclasses exhibiting a monotonically decreasing trend throughout hESC-to-CM differentiation. Values shown are global z-scores of relative molar fractions, allowing cross- stage comparison of relative abundance within each lipid subclass. Data are presented from independent biological replicates (*n* = 5–6).

Among 39 PC O subclasses, most peaked at ESC or MES stages and suddenly reduced at CPC and remained low level during CM maturation, except PC O-36:5(O-16:1_20:4), PC O-40:7(O-18:1_22:6), PC O-38:2(O-20:1_18:1), PC O-36:5, or PC O-40:7 (**Figure** 2E). Similar trends were observed for glycosylated ceramide derivatives, such as GlcCer and GalCer (**Figure** 2F, 2G).

Overall, CL, DAG, and BMP were most abundant at the ESC stage, showing relative enrichment at the pluripotent state. GalCer and GlcCer gradually decreased through early stages and sharply declined in mature CMs

### Biphasic Lipid Dynamics Peaking at Mesoderm and Cardiac Progenitor Stages During Early Differentiation

SL, characterized by a sphingoid base backbone, is an essential class of membrane components. The 14 SL subclasses identified here displayed trends largely mirroring that of Sph, with all peaking at the MES stage (**Figure** 3A, B). SL species containing hydroxylated fatty acid groups were scarce at the ESC stage but became relatively enriched at the CPC stage. In contrast, non-hydroxylated SL subtypes were abundant at ESC and MES stages. Many SL species exhibited paired molecular forms differing by hydroxylation status, indicating increased FA hydroxylation post-CM differentiation onset, reflecting stage- associated remodeling of sphingolipid hydroxylation during the MES-to-CPC transition (**Figure** 3A).

**Figure 3.**
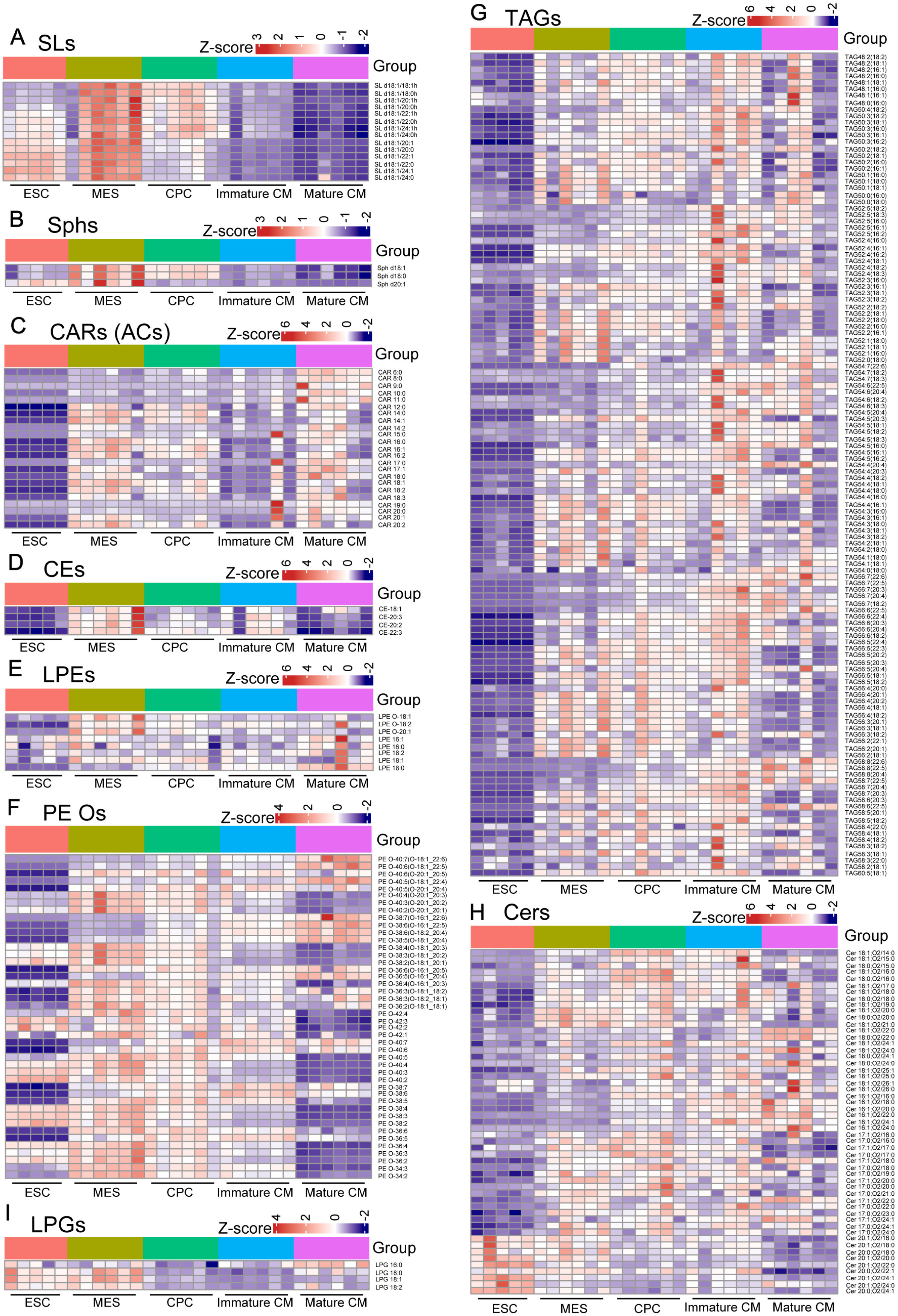
Dynamic shifts in lipid subclasses with an initial increase and subsequent decrease during hESC-derived CM differentiation (A) –(I) Heatmap presentation of dynamic remodeling patterns for all biphasic lipid subclasses that peak specifically at the mesoderm or cardiac progenitor stage and gradually decline during subsequent CM maturation. Values shown are global z-scores of relative molar fractions, allowing cross-stage comparison of relative abundance within each lipid subclass. Data are presented from independent biological replicates (*n* = 5–6).

Sph, the structural precursor in SL biosynthesis, supports the formation of major subclasses such as Cer and SM. Among the three Sph species (d18:0, d18:1, d20:1), similar temporal profiles were observed (**Figure** 3B). Levels of d18:0 and d18:1 rose sharply at the MES stage then gradually declined, reaching lowest level in mature CMs, while d20:1, initially higher at the ESC stage, peaked at the MES stage and fell markedly at the CPC stage (**Figure** 3B). These trends suggest that Sph is relatively enriched at early differentiation stages and gradually declined during maturation.

ACs (CARs), though not formally categorized as lipids, facilitate mitochondrial import of long-chain FAs for β-oxidation. Twenty-three AC species, varying from C6 to C20, were identified, displaying distinct profiles from lipid species (**Figure** 3C). Short-chain ACs (≤ C11) were markedly enriched in mature CMs, whereas longer-chain ACs reached peak abundance both during terminal differentiation and earlier stages (MES and CPC), suggesting dynamic regulation of FA metabolism across CM development (**Figure** 3C).

CEs, synthesized to neutralize free cholesterol cytotoxicity^43^, peaked during the MES stage across all four detected species, implying a role in early differentiation. Longer and more unsaturated species (CE-20:2, CE-20:3, CE-22:3) were preferentially enriched during later differentiation stages (**Figure** 3D). LPE, a PE derivative essential for mitochondrial and ER membrane synthesis, showed stage-specific distribution: the three ether-linked LPE plasmalogens (LPE O-18:1, O-18:2, O-20:1) peaked at MES, whereas the remaining species were elevated in mature CMs (**Figure** 3E).

PE O is generated via AGPS-mediated replacement of the sn-1 hydroxyl with an ether bond to a long-chain alkyl group. Of the 44 identified PE O species, 25 peaked at the MES stage, followed by a gradual decline (**Figure** 3F). Nine species were most abundant in mature CMs, whereas eight peaked at the CPC or immature CM stages. Among the 17 PE O species upregulated in late differentiation, the hydrocarbon chains typically contained five or more double bonds, with highly unsaturated species preferentially enriched at later stages, consistent with a potential contribution to membrane fluidity during maturation (**Figure** 3F).

In total, 125 TAG species were detected, with overall relative abundance peaking at the immature CM stage, independent of chain length or unsaturation degree (**Figure** 3G). Cer species displayed different dynamics. Although derivatives such as GlcCer and GalCer species mainly enriched at ESC stage (**Figure** 2F, 2G; S2B), the overall Cers reached their maximum relative abundance at the CPC stage (**Figure** 3H; S2E). LPG species were mainly enriched during the ESC and MES stages, except LPG 16:0, which peaked in Day-30 CMs, suggesting that LPG species are predominantly detected at ESC and MES stages, with only LPG 16:0 accumulating in late-stage cultures. (**Figure** 3I).

### Remodeling of Lipid Subclasses with a Down-Then-Up Pattern: Early-Stage Reduction and Late-Stage Enrichment

PS, a critical component of cellular membranes, contributes to membrane asymmetry and structural stability. Similar to SM, PS subclasses with longer acyl chains generally reach peak abundance after the CPC stage (**Figure** 4A). Among the identified 36-carbon PS species, the more unsaturated PS 36:2 (18:1_18:1) is most abundant at the human ESC stage and gradually declines thereafter, whereas the less unsaturated PS 36:1 (18:0_18:1) increases progressively during differentiation, peaking in mature CMs (**Figure** 4A). This reciprocal trend indicates a gradual increase in PS lipid saturation, accompanied by reduced acyl chain unsaturation during late CM maturation, primarily reflected in the PS 36:2 (18:1_18:1) subclass.

**Figure 4.**
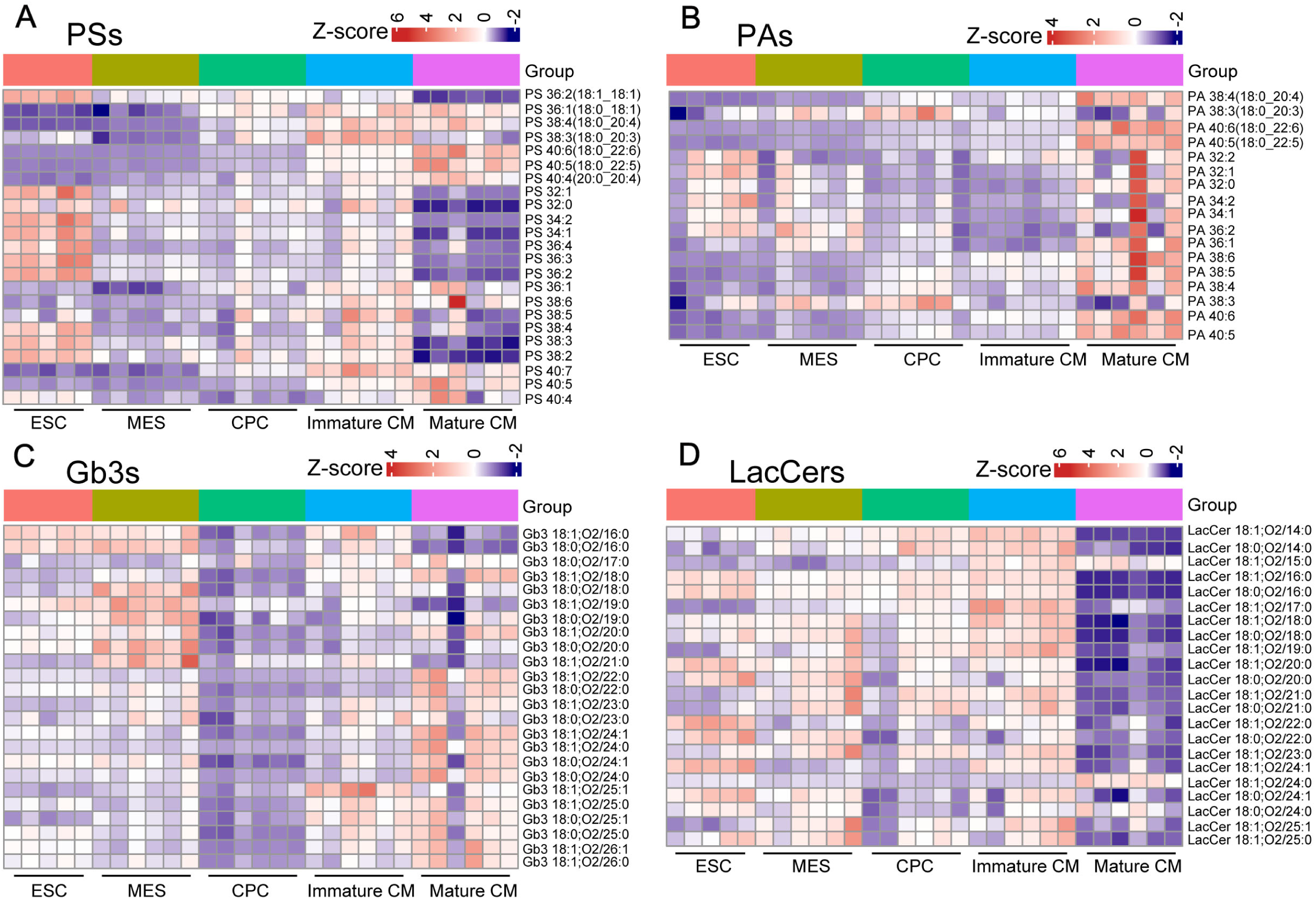
Dynamic fluctuations in lipid subclasses with an initial decrease and subsequent increase during hESC-derived CM differentiation (A) –(D) Heatmap visualization of lipid subclasses showing suppressed abundance in early differentiation stages followed by progressive elevation during late CM maturation. Values shown are global z-scores of relative molar fractions, allowing cross-stage comparison of relative abundance within each lipid subclass. Data are presented from independent biological replicates (*n* = 5–6).

Phosphatidic acid (PA), a glycerophospholipid involved in the regulation of mitochondrial dynamics and functions^44^, displayed distinct temporal lipidomic patterns (**Figure** 4B). With the exception of PA 38:3, which peaked at the CPC stage, nearly all PA species containing 38 or more carbon atoms reached maximal abundance in mature CMs. Throughout CM differentiation, PA exhibited progressive increases in acyl chain length, degree of unsaturation, and molecular complexity, consistent with the biosynthetic demands of maturing CM membranes (**Figure** 4B).

Gb3, a glycosphingolipid composed of ceramide linked to a galactose–glucose–galactose trisaccharide, was represented by 24 species. Species with longer hydrocarbon chains and more complex structures exhibited progressively later peak abundances during CM differentiation (**Figure** 4C). Gb3 levels declined markedly at the CPC stage, followed by a subsequent increase, suggesting stage-specific regulation of glycosphingolipid metabolism (**Figure** S2C).

LacCer, generated through the conjugation of ceramide and lactose, was represented by 22 subclasses. Most LacCer species peaked before the immature CM stage, whereas only LacCer 18:1;O2/24:0 achieved maximal levels in mature CMs (**Figure** 4D). Notably, GalCer and LacCer species sharply decreased from immature to mature CMs (**Figure** 4D, E; S2B, C). Overall, LacCer and GalCer are more abundant at early differentiation stages while decrease sharply in mature CMs, exhibiting a stage-specific downregulation during maturation progression of hESCs derived CMs *in vitro*.

### Progressive Accumulation of Monotonically Upregulated Lipid Subclasses Is Associated with CM Structural and Functional Maturation

FAs serve as fundamental building blocks for lipid synthesis and represent a major energy reserve in cells. Through β-oxidation, FAs are catabolized to generate ATP, particularly under fasting or high-energy-demand conditions^18–20^. In this study, 11 FA species were identified, with peak abundances occurring predominantly during the middle to late stages of CM differentiation (**Figure** 5A). Species with longer carbon chains or higher degrees of unsaturation were more enriched in later stages, reflecting the potentially increased requirement for lipid complexity and membrane fluidity in maturing CMs.

**Figure 5.**
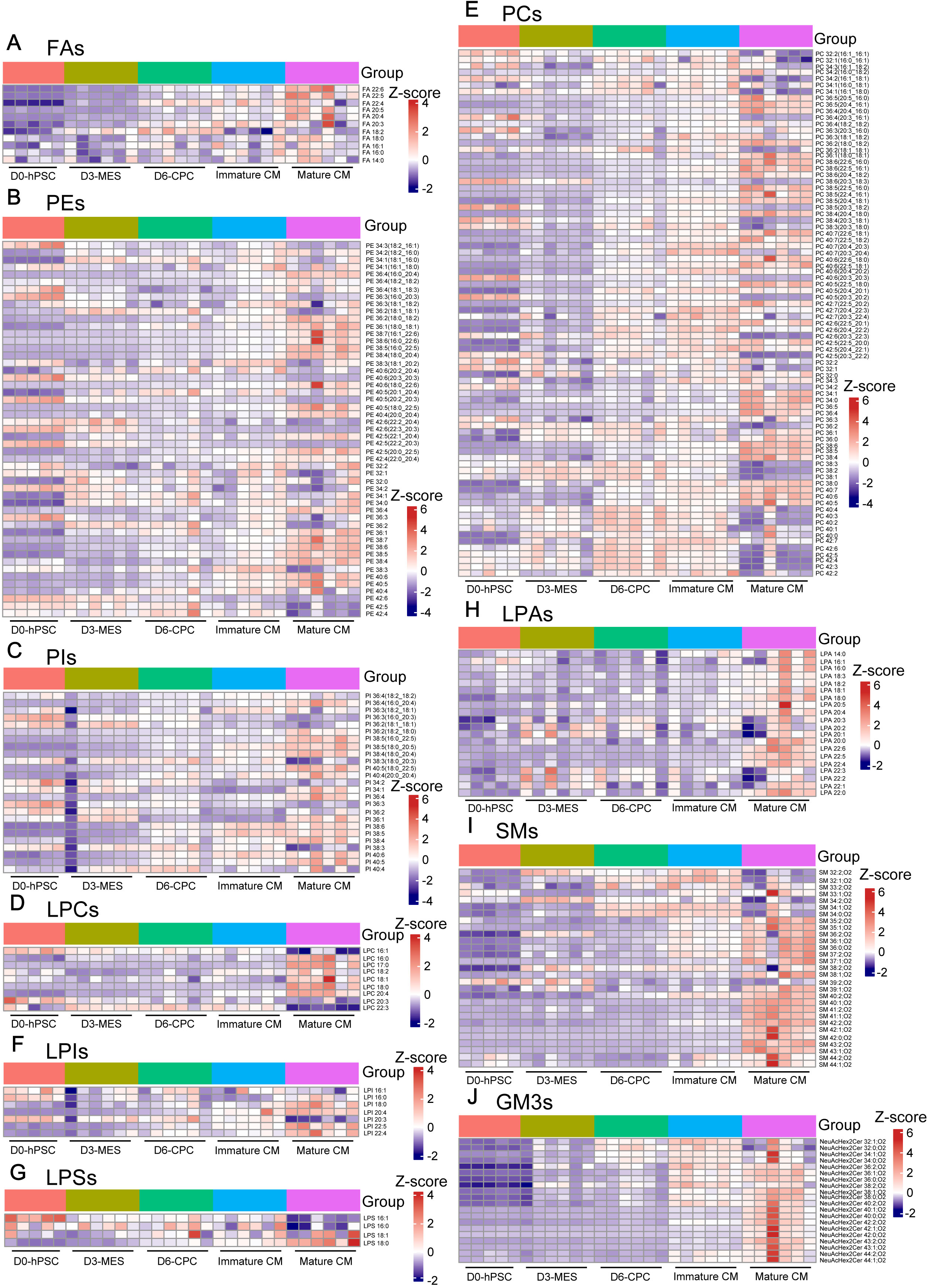
Dynamic changes in monotonically upregulated lipid subclasses during hESC-derived CM differentiation (A) –(J) Comprehensive heatmap profiling of all lipid subclasses displaying continuous, progressive accumulation across the entire CM differentiation trajectory, from hESCs to matured CMs. Values shown are global z-scores of relative molar fractions, allowing cross- stage comparison of relative abundance within each lipid subclass. Data are presented from independent biological replicates (*n* = 5–6).

PE, a major glycerophospholipid with an ethanolamine headgroup, was represented by 41 species. Among these, 23 reached maximal abundance in mature CMs, including 22 species with carbon numbers exceeding 36 and 19 with four or more double bonds (**Figure** 5B). Certain PE species, such as PE 40:5 (18:0–22:5), increased continuously throughout differentiation, whereas others, like PE 42:6 (22:3–20:3), exhibited a gradual decline. These patterns suggest progressive increases in acyl chain length, unsaturation, and structural complexity as CM differentiation advances.

PI, a key membrane phospholipid, showed a subclass-specific trend based on unsaturation degree (**Figure** 5C). Species with four or more double bonds peaked in mature CMs, while less unsaturated forms (≤ three double bonds) were predominant at earlier stages such as the ESC phase. This pattern is consistent with progressive enzymatic desaturation of phospholipid acyl chains during long-term CM culture, which has been well documented in both *in vivo* heart development and *in vitro* stem cell-derived CM maturation models, mediated by fatty acid desaturases and lysophospholipid acyltransferases such as LPCAT3 and SCD5^45–47^.

LPC (Lyso-PC), generated by phospholipase A1/A2-mediated hydrolysis of PC, plays a central role in phospholipid metabolism. Among nine detected LPC species, six peaked at the mature CM stage, independent of chain length or unsaturation (**Figure** 5D), suggesting potential importance in CM maturation and possible involvement in signaling or energy regulation.

Among the 82 PC species identified, 65 peaked during the mid-to-late stages of differentiation—namely the CPC, immature CM, and mature CM phases (**Figure** 5E). Species with 36 or more carbons and four or more double bonds predominated in mature CMs. Given their consistently high molar fraction relative to polar lipids (exceeding 0.4) across all stages, PCs likely represent a major membrane lipid component across all stages of CM differentiation. Interestingly, PC species with > 40 carbons were enriched in CPCs and immature CMs, suggesting that excessive chain elongation may reduce the necessity for high unsaturation during CM development.

LPI, a hydrolysis product of PI and an intracellular signaling molecule, exhibited subtype-specific dynamics (**Figure** 5F). Four of the seven detected LPI species, each containing > 18 carbons or ≥ four double bonds, gradually increased through differentiation and peaked in mature CMs, consistent with their regulatory roles in advanced CM stages.

LPS, represented by four subclasses, displayed chain-length–dependent accumulation during CM differentiation (**Figure** 5G). Long-chain species such as LPS 18:0 and LPS 18:1 reached peak abundance in mature CMs, whereas shorter-chain variants (LPS 16:0 and LPS 16:1) predominated early and declined thereafter.

LPA, derived from hydrolysis of PA, PC, or DAG, is crucial for cardiovascular development, including angiogenesis and cardiac morphogenesis^48,49^. Consistent with its established roles, all 20 detected LPA subspecies peaked in mature CMs (**Figure** 5H), suggesting extensive involvement in CM maturation and signaling.

SM comprised 29 subclasses, with acyl chain lengths ranging from 32 to 44 carbons and a maximum of two double bonds (**Figure** 5I). Species with ≥35 carbon atoms mainly peaked in mature CMs, indicating key contributions to CM membrane remodeling. Ganglioside GM3, a glycosphingolipid containing ceramide and sugar moieties, displayed peak levels in species with total carbon numbers ≥ 36 at the mature CM stage (**Figure** 5J). Its progressive accumulation during differentiation is consistent with a potential association with CM developmental specialization especially at later culture stages.

### Progressive Elongation and Unsaturation of Fatty Acid Chains Correlate with Membrane Adaptation During Differentiation

Lipids, especially phospholipids, form the fundamental structural components of cellular and organelle membranes. Membrane fluidity arises largely from the flexibility of phospholipids, which depends on their composition and degree of unsaturation^50–52^. To analyze lipid unsaturation, we categorized lipid molecules into three groups based on the number of double bonds in their hydrocarbon chains: saturated (no double bonds), mono- and diunsaturated (1–2 double bonds), and polyunsaturated (> 2 double bonds) lipids^26^. During human CM differentiation, lipid unsaturation levels increase alongside hydrocarbon chain elongation (**Figure** 6A). We observed that some lipids with identical chain lengths exhibit dynamic changes in unsaturation, increasing at the CPC stage but decreasing at the mature CM stage, as indicated by red arrows. Additionally, certain lipid species show elevated levels at the MES stage, as indicated by green arrows (**Figure** 6A). Quantitative analysis revealed divergent trends: mono- and diunsaturated lipids decreased sharply starting from the MES stage and continued declining, whereas polyunsaturated lipids rose markedly during MES stage before stabilizing (**Figure** 6B). Among 31 lipid categories, PCs most prominently exhibited this pattern (**Figure** 6C). Similar trends were observed relative to saturated lipids (**Figure** S3A), although in mature CMs, the polyunsaturated-to-saturated lipids ratio declined, suggesting that polyunsaturated lipids are critical throughout differentiation, while mono- and diunsaturated lipids are less essential.

**Figure 6.**
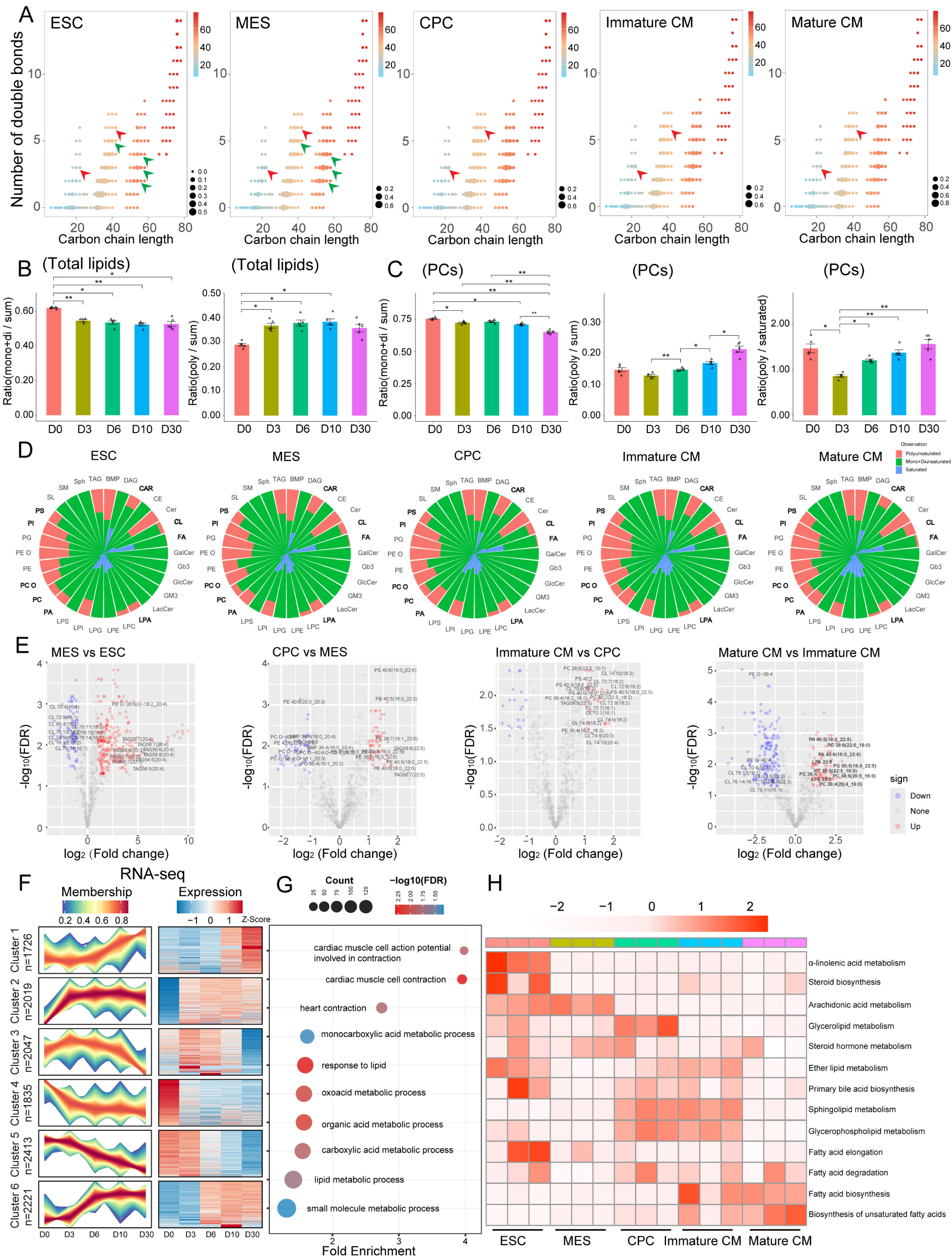
Dynamic remodeling of lipid acyl chain length and unsaturation during hESC-derived CM differentiation (A) Bubble plot summarizing global remodeling of acyl chain length and carbon-carbon double bond unsaturation degrees across sequential CM differentiation stages. The size of each bubble is proportional to the total relative molar fraction of lipids with corresponding chain- length and unsaturation characteristics. Red arrows denote representative lipid species with stage-dependent increasing relative abundance; green arrows indicate species with declining temporal trends. (B) Quantitative bar statistics comparing molar ratio percentages of combined mono- plus diunsaturated lipids and polyunsaturated lipids relative to the total polar lipid pool across five differentiation stages. Each dot corresponds to one independent biological replicate (*n* = 5). Data are presented as mean ± SEM. Overall inter-stage statistical differences were determined using one-way ANOVA followed by Tukey’s post-hoc test. (C) Stagewise ratios of unsaturated species within the phosphatidylcholine (PC) lipid family, including mono- plus diunsaturated PC/total PC and polyunsaturated PC/saturated PC ratios across developmental time points. Individual dots represent biological replicates (*n* = 5); data are shown as mean ± SEM. Statistical evaluation among multiple staged groups was performed using one-way ANOVA followed by Tukey’s post-hoc test. (D) Nightingale rose charts illustrating dynamic proportional redistribution of saturated, mono + diunsaturated, and polyunsaturated lipid species across the full CM differentiation time course. (E) Volcano plots showing significantly differentially altered lipid species across all four consecutive developmental transitions: ESC vs MES, MES vs CPC, CPC vs immature CMs, and immature CMs vs mature CMs. The most prominent lipid remodeling is observed during the late-stage extended culture transition from immature to mature CMs. Red dots denote statistically significantly upregulated lipid species; blue dots indicate significantly downregulated species. Significance thresholds were defined as |log₂(fold change)| > 1.0 and BH-FDR adjusted *p* < 0.01. Statistical testing was performed using two-tailed unpaired Student’s *t*-test followed by BH-FDR correction for multiple testing. (F) Time-series clustering of transcriptomic profiles of all dynamically expressed genes based on temporal expression trajectories across CM differentiation stages using the Mfuzz soft clustering algorithm. Membership values range from 0 to 1 (dimensionless), and expression levels are shown as z-scores (dimensionless; relative expression in standard deviation units). (G) Functional GO enrichment analysis specifically for genes in Cluster 1, utilizing an expressed-gene background (mean TPM>1.0 in at least one stage). Representative terms with BH-FDR adjusted *p* < 0.05 are displayed. (H) Heatmap of pathway-level scores across stages. Pathway scores represent the mean expression levels (TPM) of all genes assigned to each respective pathway. Statistical significance of dynamic regulation across stages was determined by one-way ANOVA with BH-FDR correction. Detailed pathway lists and statistical results are provided in Supplementary Table S4. Data are presented from independent biological replicates (*n* = 3).

Subclass analyses highlighted increases in polyunsaturated species of PC, CAR, CL, FA, LPA, PA, PC O, PI, and PS (**Figure** 6D). Notably, CE and PA polyunsaturated subclasses slightly decreased at the mature CM stage, whereas PC increased. Polyunsaturated PE transiently decreased at MES stage before rising. Saturated lysophospholipids, including LPC, LPE, LPG, and LPS, were increased, whereas polyunsaturated LPC species were decreased (**Figure** 6D).

Volcano plot analysis across all consecutive developmental stages revealed dynamic lipid remodeling throughout the entire differentiation trajectory, with the most pronounced changes occurring between immature and mature CMs. Specifically, selected individual species within the PC, PA, and LPA classes showed the most prominent increases at the mature stage (**Figure** 6E). Full pairwise comparisons across all non-consecutive stages are provided as supplemental data in **Figure** S3B. Mfuzz clustering of RNA-seq expression trajectories revealed that genes in Cluster 1 exhibited an upregulation trend towards later stages of differentiation (**Figure** 6F). Rigorous GO enrichment analysis of Cluster 1 (using an expressed-gene background) highlighted the enrichment of terms essential for CM maturation, including both functional processes (e.g., cardiac muscle cell contraction and action potential) and key metabolic transitions (e.g., lipid, carboxylic acid, and oxoacid metabolic processes) (**Figure** 6G, **Supplementary Table** S1). Furthermore, quantitative pathway-level analysis confirmed that fatty-acid and lipid metabolic pathways, including the biosynthesis of unsaturated fatty acids, are dynamically regulated during differentiation, showing significant upregulation in later stages of CM maturation (Day-10 and Day-30 CMs) (**Figure** 6H, **Supplementary Table** S4).

These data collectively indicate that during human embryonic stem cell differentiation into CMs, there is a progressive increase in polyunsaturated lipids. This likely reflects functional requirements for enhanced membrane fluidity, which is critical for effective CM contractility.

### Correlation Between PC Metabolism Dynamics and Transcriptional Reprogramming During CM Differentiation and Maturation

Referring to ATAC-seq data from different stages of CM differentiation reported previously^53^, we focused on the regulatory regions of genes involved in the biosynthesis of unsaturated fatty acids. By performing motif scanning on these specific peak regions, we identified a predicted transcription factor (TF) network (**Figure** 7A, S4A). This network highlights candidate TFs, including SP1, KLF4, IRF1, IRF2, USF1, FOXO1, IRF4, and CREB1, which are predicted to be associated with fatty acid synthesis and cardiac development^54–56^. To identify genes synergistically regulated at both the epigenetic and transcriptional levels during late-stage maturation, we intersected ATAC-seq Cluster 5 (representing regions with progressively increasing chromatin accessibility) with RNA-seq Cluster 1 (representing genes with progressively increasing expression) (**Figure** 7B, **Supplementary Table** S5). This strict intersection yielded 376 overlapping genes. Rigorous Gene Ontology (GO) enrichment analysis of these 376 genes, utilizing an expressed-gene background and strictly filtering for FDR < 0.05, highlighted critical pathways predominantly involved in extracellular matrix (ECM) organization, cell communication, and structural morphogenesis (**Figure** 7C, **Supplementary Table** S5). This indicates that the ultimate phenotypic output of the epigenetic and transcriptional reprogramming during late-stage maturation is the structural and functional integration of the myocardial tissue. While the global multi-omics integration highlighted a continuous upregulation of structural and communicative pathways towards Day 30, we observed a distinct dynamic pattern when specifically examining targeted lipid metabolic genes. Promoter-region accessibility heatmaps for selected genes associated with PC, PA, LPC, and LPA metabolism (**Figure** 7D-G; **Supplementary Table** S6) revealed that their chromatin accessibility generally peaked at the CPC stage and subsequently decreased by Day 30. We note that these panels represent descriptive associations rather than establishing causal regulation of lipid biosynthesis. Transcriptomic analysis of lipid synthesis-related genes showed convergent overall expression trends across various differentiation stages (**Figure** S4B-F). In mature CMs, the biosynthesis of PC, PA, LPC/LPA, and GM3 is predominantly mediated by specific enzymes such as CHPT1, PLD1, PLA2G1B, and UGCG/ST3GAL5, respectively. Given the complexity of lipid metabolism, individual genes can regulate multiple lipid species. Ultimately, the contrast between the continuous structural demands of the maturing tissue (**Figure** 7C) and the transient epigenetic priming followed by a dampening of endogenous lipid metabolic accessibility (**Figure** 7D-G) led us to hypothesize that this endogenous metabolic shift might render the maturing CMs responsive to exogenous lipid supplementation. Therefore, we next investigated whether exogenous PC supplementation could further modulate the transcriptomic and structural state of differentiating CMs.

**Figure 7.**
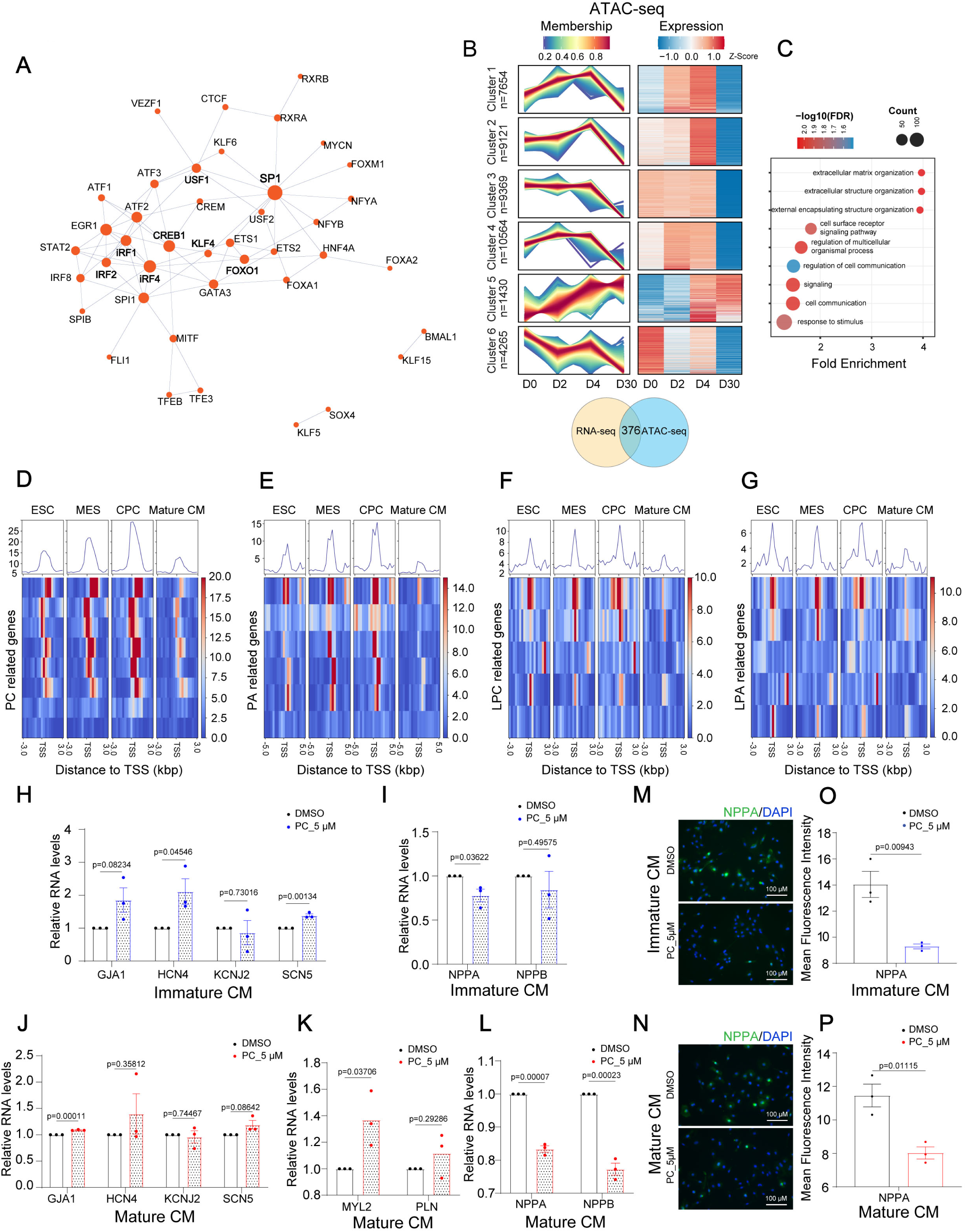
Altered PC metabolism correlates with hESC-derived CM differentiation and maturation progression (A) Predicted transcriptional regulatory network specific to lipid metabolism. The network was generated by performing motif scanning (using the SEA tool) on ATAC-seq peaks^53^ located in the regulatory regions of genes involved in the Biosynthesis of Unsaturated Fatty Acids pathway, integrated with in-house temporal RNA-seq data. Edges represent predicted regulatory relationships based on motif enrichment. Candidate core transcription factors are highlighted. (B) Mfuzz clustering of dynamic genome-wide chromatin accessibility peaks across CM differentiation stages. ATAC-seq peaks were assigned to target genes based on proximity. The Venn diagram illustrates the strict intersection yielding 376 overlapping genes between RNA-seq expression Cluster 1 (progressively increasing expression) and ATAC-seq Cluster 5 (progressively increasing accessibility). Membership values range from 0 to 1 (dimensionless), and expression levels are shown as Z-scores (dimensionless; relative expression in standard deviation units). (C) GO functional enrichment analysis of the 376 overlapping target genes identified in (B). The analysis utilized an expressed-gene background (mean TPM ≥ 1.0 in at least one stage), and only terms with BH-FDR-adjusted p < 0.05 are displayed. (D) –(G) Temporal heatmap visualization of chromatin accessibility signal dynamics at promoter regions of selected lipid metabolic genes associated with PC (D), PA (E), LPC (F), and LPA (G) metabolism throughout CM differentiation. (H) Relative mRNA expression levels of electrophysiology and CM contractility-associated genes in immature CMs following 13-day continuous culture with 5 μM PC supplementation versus DMSO vehicle control group. All data are presented as mean ± SEM from independent biological replicates (*n* = 3). Inter-group differences were compared using two- tailed unpaired Student’s *t*-test with BH-FDR correction for multiple gene comparisons. (I) Relative transcript levels of cardiac hypertrophy marker genes in immature CMs under PC supplementation versus control conditions. All data are presented as mean ± SEM from independent biological replicates (*n* = 3). Statistical comparison was performed using two- tailed unpaired Student’s *t*-test with BH-FDR correction for multiple gene comparisons. (J) Quantitative comparison of CM functional maturation gene expression in mature CMs after prolonged PC intervention relative to vehicle control. All data are presented as mean ± SEM from independent biological replicates (*n* = 3). Significance analysis was conducted using two-tailed unpaired Student’s *t*-test with BH-FDR correction for multiple gene comparisons. (K) Transcriptional levels of canonical CM developmental markers in PC-supplemented versus control mature CMs. All data are presented as mean ± SEM from independent biological replicates (*n* = 3). Group differences were assessed using two-tailed unpaired Student’s *t*-test with BH-FDR correction for multiple gene comparisons. (L) Hypertrophy-related gene expression profiling in mature CMs with or without PC supplementation. All data are presented as mean ± SEM from independent biological replicates (*n* = 3). Two-tailed unpaired Student’s *t*-test was used for statistical comparison between two groups with BH-FDR correction for multiple gene comparisons. (M, O) Quantitative IF intensity analysis of NPPA protein expression in PC-treated and DMSO-treated immature CMs. Scale bar, 100 µm. All data are presented as mean ± SEM from independent biological replicates (*n* = 3). Statistical comparison was performed using two-tailed unpaired Student’s *t*-test. (N, P) Quantified NPPA IF signal intensity in mature CMs with or without PC metabolic intervention. Scale bar, 100 µm. All data are presented as mean ± SEM from independent biological replicates (*n* = 3). Two-tailed unpaired Student’s *t*-test was applied for statistical evaluation.

Compared to controls, supplementation with 5 μM PC for 13 days was accompanied by altered expression levels of genes associated with electrophysiological regulation in immature CMs (day 20), including HCN4, a core heart rate modulator, and SCN5A, encoding the cardiac sodium channel responsible for myocardial action potential depolarization (**Figure** 7H). At this stage, myocardial identity markers showed no significant differences (**Figure** S4G), although hypertrophy-associated gene such as NPPA (atrial natriuretic peptide) was modestly downregulated (**Figure** 7I). In mature CMs, PC supplementation moderately increased expression of GJA1, encoding gap junction proteins critical for electrical signal propagation between cardiac muscle cells (**Figure** 7J), and MYL2, a myocardial maturity marker (**Figure** 7K). Concurrently, hypertrophy-related genes NPPA and NPPB (B-type natriuretic peptide) were significantly downregulated (**Figure** 7L). IF analysis of NPPA confirmed these transcriptional changes (**Figure** 7M, O; N, P). Collectively, these results suggest that PC supplementation is correlated with elevated transcriptional levels of partial mature CM marker genes and improves the CM viability under experimental conditions, indicating its therapeutic potential for hypertrophy-related cardiac diseases. However, the functional consequences of these changes require further investigation.

## DISCUSSION

Metabolic remodeling during heart injury and development involves substantial shifts in energy substrate utilization and mitochondrial function^57^. In hyperlipidemia-induced fibrosis, PPARγ-driven lipid remodeling accelerates progression^58^. In diabetes, epicardial adipose tissue shows dysregulated lipid secretion, marked by saturated non-esterified fatty acids (FAs) and specific ceramides (Cer16:0, Cer24:1)^59^. Diabetes also impairs endothelial–CM communication, with cardiac endothelial cells exhibiting aberrant long-chain acylcarnitine accumulation^60^. By contrast, perivascular adipose tissue maintains vascular homeostasis via polyunsaturated FA-derived oxylipins, including linoleic acid, arachidonic acid, and epoxyeicosatrienoic acids (EETs)^61^. However, no systematic study has yet analyzed lipidomic changes throughout CM differentiation. Using low-injection-volume ultra-sensitive targeted metabolomics and lipidomics^26^, we comprehensively profiled distinct lipid class remodeling during *in vitro* CM differentiation from ESCs (**Figure** 1), defining the dynamic lipidome of this process. We identified 781 lipid species across 31 major classes exhibiting five distinct temporal trends. Notably, CL, PE, and PC classes showed significant increases in acyl chain length and unsaturation (**Figure** 2A, 5B and 5E). Concurrently, CL acyl chains shifted from C16:1 toward C18:2, C20:3, and C20:4. Co-regulation analysis among CL, PC, and PE suggested that during *in vitro* CM differentiation, PE may supply polyunsaturated acyl substrates for CL remodeling, or PE and CL may act cooperatively during metabolic transition.

In this study, we performed multi-dimensional validation of Day-30 CMs, including sarcomeric structure (ACTN2), mitochondrial morphology (TIM23), and electrophysiological properties (MEA). The observed increases in sarcomere length, mitochondrial elongation and sarcomere-aligned distribution, and prolonged field potential duration are all consistent with well-established trajectories of CM maturation progression^22,23^. These validations support that the lipidomic remodeling observed here is associated with the acquisition of maturation-associated features in the *in vitro* CM culture system.

Based on the observed stage-specific abundance patterns, we propose several testable hypotheses regarding the roles of lipids during CM differentiation. First, the elevated levels of CL, DAG, BMP, and simple glycosphingolipids at the pluripotent stage may support rapid membrane biogenesis and maintenance of stem cell identity; targeted perturbation of CL or glycosphingolipid biosynthesis will be necessary to validate their roles in lineage priming. Second, the transient enrichment of sphingolipids, acylcarnitines, and ether phospholipids at the mesoderm and cardiac progenitor stages suggests that these lipid classes may contribute to mesodermal fate transitions and progenitor state maintenance. Third, the progressive accumulation of PC, LPA, SM, and GM3 species at later stages is consistent with potential roles in membrane remodeling and CM maturation. However, these interpretations remain correlative and will require functional validation through targeted metabolic perturbation.

Lipidomics revealed that polyunsaturated lipid content increased significantly after CM differentiation onset, while mono- and diunsaturated lipids decreased, changes persisting through subsequent stages. These results highlight distinct stage-specific lipidomic features across the full course of hESCs-derived CM differentiation *in vitro*.

We hypothesize that the progressive increase in polyunsaturated phospholipids enhances membrane fluidity and dynamic responsiveness, thereby accommodating the increased contractile activity of late-stage CMs. This hypothesis is testable and will require direct measurements of membrane biophysical properties alongside functional assessment of contractility in future studies.

Integrated ATAC-seq and RNA-seq analyses showed significant gene expression changes related to unsaturated fatty acid synthesis, biological processes, and signaling pathways that matched lipidomic profiles (**Figure** 6, 7). Genes with consistent RNA-seq upregulation were enriched in anabolic pathways including FA, long-chain FA, PC, and TAG synthesis. Genes upregulated in both ATAC-seq and RNA-seq were enriched in TAG metabolism, cardiac muscle development, and electrical conduction pathways, consistent with lipidomic observations of increased lipid unsaturation, chain length, and changes in PC, TAG, and FA. These transcriptional patterns are consistent with the observed lipidomic remodeling of acyl chain unsaturation and length, and may underpin cellular adaptations including membrane property changes, metabolic transition, and contractile function in late-stage CMs. Notably, changes in major lipid classes do not fully correlate with their subclasses, likely due to varying cellular lipid subclass content.

Our data show that lipid species with progressive accumulation during differentiation, including PC, LPC, PA, and LPA, are enriched at mature CM stages. Myocardial hypertrophy comprises physiological (PhCH) and pathological subtypes^62^. PhCH occurs during postnatal growth, exercise, or specific conditions, characterized by adaptive CM enlargement and is crucial for cardiac development^63^. Aberrant expression of hypertrophy-related genes outside normal temporal regulation can impair cardiac development.

Given that physiological hypertrophy is a normal component of CM developmental progression, the observed downregulation of selected hypertrophy-associated transcripts following PC supplementation suggests a potential association between phospholipid availability and CM developmental state. We propose that exogenous PC supply may modulate hypertrophic gene programs from CPC stage; however, this study only assessed transcript-level changes, and whether these effects translate to functional improvements in maturation, contractility, or stress resistance remains to be determined. The predicted transcription factor network derived from multi-omics integration provides candidate regulators of lipid metabolic remodeling, but direct evidence of regulatory causality requires further chromatin immunoprecipitation and functional perturbation assays.

We hypothesize that simple lipid species in early-stage ESCs support rapid membrane assembly and biosynthetic precursor supply, whereas complex lipids in later differentiation stages contribute to membrane stability, fluidity during contractile activity, and intracellular second messenger signaling. Lipidome remodeling partially reflects the metabolic shift from glycolysis-dominated stem cells to matured CMs reliant on oxidative phosphorylation.

### Limitations of the study

First, although Day-30 CMs exhibit core structural, mitochondrial, and electrophysiological maturation-associated features, they are generated via standard *in vitro* monolayer culture and do not fully recapitulate the complete structural, metabolic, and functional maturity of adult *in vivo* CM s. These cells represent a relatively advanced maturation stage within the *in vitro* differentiation system, and the observed lipid remodeling reflects *in vitro* culture-associated developmental progression. Future studies using dedicated maturation-promoting strategies (e.g., 3D culture, mechanical stretch, fatty acid conditioning) and primary adult CMs are needed to extend these findings.

Second, lipidomic profiling was performed on unsorted bulk culture lysates. Although quantitative validation confirmed that CMs account for > 88% of the population from day 10 onward, a small proportion of non-myocyte cells remains as expected in standard directed differentiation. The observed lipid patterns primarily reflect CM -intrinsic remodeling, but potential contributions from non-myocyte populations are acknowledged; future studies using purified CM populations would help confirm cell-intrinsic lipid dynamics.

Third, the lipidomics data represent relative molar fractions within the measured polar lipid pool, not absolute abundance per cell. Compositional data supports correlative interpretations, and future absolute quantification and functional perturbation experiments are needed to establish causal roles of specific lipid species in CM developmental progression. Additionally, standard statistical tests applied to compositional molar fraction data have inherent limitations due to the non-independence of lipid classes, and all interpretations are correspondingly cautious.

Finally, the PC supplementation experiments only assessed transcript-level and translational level changes; functional validation at the electrophysiological, metabolic, and contractile levels is required to determine the physiological consequences of altered phospholipid availability.

Given the exploratory nature of current research, future studies should investigate the regulatory genes linked to lipid unsaturation and specific lipid changes during cardiac differentiation to elucidate their regulatory mechanisms and physiological roles in greater depth.

## RESOURCE AVAILABILITY

### Lead Contact

Further information and requests for resources and reagents should be directed to and will be fulfilled by the lead contact, Wuming Wang.

### Materials Availability

This study did not generate new unique reagents.

### Data and code availability

Bulk RNA-seq raw data and normalized mapped reads have been deposited at GEO (GSE310576). Lipid metabolism datasets have been deposited at Mendeley Data (DOI: 10.17632/r5db9w567v.2). All original code has been deposited at a Zenodo repository (DOI: https://doi.org/10.5281/zenodo.20102571). DOIs are listed in the **KEY RESOURCE TABLE**. Any additional information required to reanalyse the data reported in this paper is available from the lead contact upon request.

## ACKNOWLEDGEMENTS

This work was supported by a Natural Science Foundation of China (NSFC)-Young Scientists Fund (#82200314), Innovation and Technology Commission of the Hong Kong Government grants (GHX09721SZ and MHP00523), Direct Grant for Research (#2021.069 and #2022.082) from CUHK, the National Natural Science Foundation of China (NSFC)/Research Grants Council (RGC) Joint Research Scheme (#N_CUHK428/22), and Strategic Seed Funding for Collaborative Research Scheme (#MK/WW/SSFCRS2324/8014/24jh) from CUHK, and RGC general research fund (#14116825). This work was partially supported by Science and Technology Planning Project of Guangdong Province, China (2023B1212120009), CUHK Laboratory Support Special Fund for MOE Key Laboratory for Regenerative Medicine (2622009), Co-funding Mechanism on Joint Laboratories with the CAS (JLFS/M-403/24), the Chinese University of Hong Kong (CUHK)–Shandong University (SDU) Joint Laboratory on Reproductive Genetics, Hong Kong Branch of CAS Center for Excellence in Animal Evolution and Genetics (8601010 and 8601011), and the research fund to the Center for Neuromusculoskeletal Restorative Medicine from Health@InnoHK program launched by Innovation and Technology Commission, Hong Kong.

## AUTHOR CONTRIBUTIONS

W.M.W. and W.Y.C conceived and designed the research; H.T.T., R.L., and Q.Q.H. performed the experiments; Z.Q.X. and G.L. conducted bioinformatic analyses; H.T.T., R.L., and W.M.W. drafted the original manuscript; H.T.T., Z.Q.X., and W.M.W. revised the manuscript and performed formal editing; W.Y.C. and W.M.W. acquired funding and supervised the project.

## DECLARATION OF INTERESTS

The authors declare no conflict of interest.

## STAR★METHODS

### Experimental model and study participant details

This study used the established human embryonic stem cell line (H1/WA01) obtained from WiCell. The original derivation of this line was ethically approved (NIHhESC-10-0043). No additional ethics approval was required for *in vitro* work with this established hESC line.

All cells were maintained in a humidified incubator at 37 °C with 5% CO₂. hESCs were cultured on Matrigel-coated culture plates in mTeSR1 medium. Cardiomyocyte differentiation was induced via sequential small-molecule modulation of WNT signaling in CM culture medium (CMCM), consisting of RPMI 1640 (Gibco, 11875085) supplemented with B27 minus insulin (Gibco, A1895601). Five consecutive developmental stages were profiled across the study: embryonic stem cells (ESC, Day 0), mesoderm (MES, Day 3), cardiac progenitor cells (CPC, Day 6), immature CMs (Day-10 CM), and mature CMs (Day- 30 CM).

This study used the established human embryonic stem cell line (H1/WA01) obtained from WiCell. The original derivation of this line was ethically approved and donors provided informed consent (WiCell). In accordance with The Chinese University of Hong Kong’s research governance, no additional ethics approval was required for *in vitro* work with this established hESC line.

## Method details

### Cell culture and passaging

H1 ESCs were cultured in a 37 ℃ incubator with 5% CO₂ using mTeSR1 medium. For each cell passage, culture plates were pre-coated with Matrigel for 1 hour, and mTeSR1 medium was supplemented with 10 µM Y-27632 2HCl (Selleckchem, S1049) during the process. H1 ESCs were dissociated with Accutase reagent at 37 ℃ for approximately 5 minutes before replating.

CMs derived from H1 ESCs were cultured on 6-well cell culture plates or 6 cm diameter cell culture dishes precoated with Matrigel (Corning, 354277) for 1 hour at 37 ℃. After approximately 10 minutes of dissociation using Cardiomyocyte Dissociation Kit (Stemcell, 05025) according to the instructions, the CMs were replated onto a new culture dish.

The study was conducted using the male H1 human ESCs. Because female cell lines were not included, whether biological sex influences cardiac differentiation, lipid remodeling, or cardiac maturation could not be determined. This may limit the generalizability of the findings across biological sexes. Pluripotency marker expression, including OCT4, SOX2, and NANOG, has been examined by immunofluorescence (IF) staining and qPCR to confirm undifferentiated status. Regular tests for mycoplasma contamination using the Myco- Blue^®^ Mycoplasma Detector kit (Vazyme) were negative.

### CM differentiation

CM differentiation was initiated by activating WNT signaling with 6 μM CHIR99021 (Selleckchem, S1263) diluted in CMCM for 48 hours, followed by culture in CMCM alone for 24 hours; this sequential treatment induced progression to the CM stage. Subsequent treatment with 5 μM IWR-1-endo (Selleckchem, S7086) diluted in CMCM for another 48 hours resulted in the formation of CPCs. Continuous culture with CMCM for approximately 3 days induced the transformation of CPCs into immature CMs, while mature CMs were generated by day 30 after the initiation of differentiation. CMs were collected at different stages according to experimental requirements and stored in methanol (Honeywell, 3221) at -80 ℃ for subsequent lipid extraction.

### IF staining

The cells were replated into new culture plates and incubated for 12-24 hours before treatment. They were then exposed to 0.15% Triton X-100 (USB, 9002-93-1) combined with 2% paraformaldehyde (Sigma, 30525-89-4) for approximately 20 minutes, followed by blocking with 5% BSA. Subsequently, cells were incubated overnight at 4 ℃ with primary antibodies: anti-NPPA (PTG 27426-1-AP), anti-SOX17 (R&D Systems, AF1924), anti-GATA4 (Cell Signaling Technology, 36966), anti-NKX2.5 (Cell Signaling Technology, 8792), anti- cTNT (Invitrogen, MA5-12960), anti-ACTN2 (Sigma, A7811), anti- TIM23 (Cell Signaling Technology, 34822), and anti-MLC2v (Proteintech, PTG10906-1-AP). After washing, secondary antibodies—goat anti-rabbit 555 (A21428), goat anti-rabbit 488 (A11008), and goat anti-mouse 555 (A21424) from Invitrogen—were applied for 1-2 hours at room temperature. DAPI (Sigma, D9542) was used for nuclear staining. Imaging was performed using a Ti-2 inverted fluorescence microscope (Nikon).

### Mitochondria, Sarcomere and MLC2v IF Co-Staining and Quantification

To assess sarcomeric organization and structural maturation of cardiomyocytes (CMs), IF co-staining was performed using a modified protocol. Cells were fixed with 2% paraformaldehyde for 20 min, permeabilized with 0.15% Triton X-100 for 15 min, and blocked with 5% BSA for 1 h at room temperature. Samples were then incubated overnight at 4 °C with a primary antibody cocktail comprising mouse anti-ACTN2 (Sigma, A7811; Z-line marker), rabbit anti-TIM23 (Cell Signaling Technology, 34822; mitochondrial marker), and rabbit anti-MLC2v (Proteintech, 10906-1-AP; ventricular A-band marker). After washing, cells were incubated with Alexa Fluor 488-conjugated goat anti-mouse IgG and Alexa Fluor 555- conjugated goat anti-rabbit IgG (Invitrogen) for 1 h at room temperature, followed by DAPI nuclear staining. Images were acquired using a Zeiss LSM 880 confocal microscope. Sarcomere length was quantified in ImageJ by measuring the distance between adjacent α- actinin-positive Z-lines in well-aligned myofibrils.

### Microelectrode Array (MEA) Electrophysiological Recording

Spontaneous electrophysiological activity of CMs was recorded using a multi-electrode array (MEA) system (Maxwell Biosystems). Day 30 CMs were dissociated using the Cardiomyocyte Dissociation Kit (STEMCELL Technologies, 05025) as described above and replated at equal density onto Matrigel-precoated MEA chips. After a 3-day recovery period to allow cell attachment and synchronization of beating, wells exhibiting stable spontaneous activity were selected for analysis. Field potentials were recorded for 5 min per well at 37 °C under 5% CO₂. Electrophysiological parameters were subsequently quantified.

### RNA isolation and quantitative real-time polymerase chain reaction

Total RNA was extracted from cultured cells or frozen tissue samples using RNAiso Plus reagent (Takara, 9109) according to the manufacturer’s protocol, and cDNA was generated according to the protocol of PrimeScript™ RT Master Mix (Takara, RR036A) subsequently. qPCR was performed according to the reaction system provided by SYBR Green Universal Master Mix (ABI, 4309155) on an ABI QuantStudio 7 Real-Time PCR System (Thermo Fisher Scientific).

### Bulk RNA-seq

Bulk RNA-seq was performed using the BGISEQ-500 platform (101-bp paired-end reads) with a target depth of 20–30 million reads per sample on average. Raw RNA-seq reads were initially preprocessed through adapter trimming and quality filtering using Trimmomatic. Subsequently, the filtered reads were aligned to the human reference genome (GRCh38) with STAR. Temporal gene expression patterns were analyzed using the R package Mfuzz. Functional enrichment analysis of Gene Ontology terms was conducted via the DAVID database (https://davidbioinformatics.nih.gov/). For quantitative pathway-level analysis (Figure 6H), pathway scores were calculated as the mean expression levels (TPM) of all genes assigned to each respective pathway. Gene lists for these specific metabolic pathways were primarily curated from the KEGG database.

### Bulk ATAC-seq

We re-analyzed previously published ATAC-seq data (GEO: GSE85330)^53^. Raw read quality was assessed, adapters were trimmed, and low-quality sequences were filtered using fastp. Filtered reads were aligned to the reference genome with Bowtie2. Genome track files (e.g., BigWig) were generated for visualization, and accessible chromatin regions were identified using MACS2 peak calling. To characterize dynamic changes in chromatin accessibility across developmental stages, we merged peaks identified from all samples to create a consensus peak set and quantified accessibility signals across all time points for these regions. We then performed Mfuzz clustering to identify clusters with specific temporal patterns.

To integrate ATAC-seq and RNA-seq data and identify genes synergistically regulated during late-stage maturation, we intersected ATAC-seq Cluster 5 (representing regions with progressively increasing chromatin accessibility, assigned to 4,284 genes based on promoter/enhancer proximity) with RNA-seq Cluster 1 (representing genes with progressively increasing expression, n=1,726 genes). This strict intersection yielded 376 overlapping genes.

To identify key transcription factors (TFs) regulating unsaturated FA biosynthesis, we first identified consensus ATAC-seq peaks located within promoter and enhancer regions of genes associated with this pathway. Motif scanning was then performed on these specific peaks using the SEA (Simple Enrichment Analysis) tool from the MEME Suite to predict potential TFs, and the resulting regulatory network was visualized using Cytoscape. To prioritize key TFs, motif-based predictions were integrated with RNA-seq expression profiles. TFs were prioritized if they exhibited either a continuous increase in expression or consistently high expression levels across developmental stages. Additionally, TSS and flanking regions accessibility heatmaps were generated for selected lipid metabolic genes (PC, PA, LPC, LPA) to illustrate descriptive associations between epigenetic states and transcriptional reprogramming.

### Lipid extraction

Lipids were extracted from intact cells using a modified Bligh and Dyer method as described previously^64^. Briefly, cells were homogenized in 750 µL of chloroform: methanol: MilliQ H_2_O (3:6:1) (v/v/v). The homogenate was then incubated at 1500 rpm for 1 h at 4 ℃. At the end of the incubation, 350 µL of deionized water and 250 µL of chloroform were added to induce phase separation. The samples were then centrifuged and the lower organic phase containing lipids was extracted into a clean tube. Lipid extraction was repeated once by adding 450 µL of chloroform to the remaining cells in aqueous phase, and the lipid extracts were pooled into a single tube and dried in the SpeedVac under organic mode. Samples were stored at -80 ℃ until further analysis.

### Lipidomics analyses

Lipidomic profiling was performed by Changzhou Lipidall Biotechnology Co., Ltd. (Jiangsu, China) using an ExionLC-AD coupled with Sciex QTRAP 7500 PLUS, as reported previously^65^. Separation of individual lipid classes of polar lipids by normal phase (NP)- HPLC was carried out using a TUP-HB silica column (i.d. 150x2.1 mm, 3 µm) with the following conditions: mobile phase A (chloroform: methanol : ammonium hydroxide, 89.5:10:0.5) and mobile phase B (chloroform : methanol : ammonium hydroxide : water, 55:39:0.5:5.5). Each of the 31 lipid categories was paired with a corresponding stable isotope internal standard.

Additionally, changes in the response intensity of these stable isotope internal standards can be used to monitor the stability of the instrument system. MRM transitions were set up for comparative analysis of various polar lipids. Individual lipid species were quantified by referring to spiked internal standards. d9-PC32:0(16:0/16:0), d9-PC36:1p(18:0p/18:1), d7- PE33:1(15:0/18:1), d9-PE36:1p(18:0p/18:1), d31-PS(d31-16:0/18:1), d7-PA33:1(15:0/18:1), d7-PG33:1(15:0/18:1), d7-PI33:1(15:0/18:1), C17-SL,d5-CL72:8(18:2)4, Cer d18:1/15:0-d7, d9-SM d18:1/18:1, C8-GlcCer, C8-GalCer, d3-LacCer d18:1/16:0, Gb3 d18:1/17:0, d7- LPC18:1, d7-LPE18:1, C17-LPI, C17-LPA, C17-LPS, C17-LPG, d17:1 Sph, were obtained from Avanti Polar Lipids. GM3-d18:1/18:0-d_3_ was purchased from Matreya LLC. Free fatty acids were quantified using d_31_-16:0 (Sigma-Aldrich) and d_8_-20:4 (Cayman Chemicals).

Glycerol lipids including DAGs and TAGs were quantified using a modified version of reverse phase HPLC/MRM^66^. Separation of neutral lipids was achieved on a Phenomenex Kinetex-C18 column (i.d. 4.6x100 mm, 2.6 µm) using an isocratic mobile phase containing chloroform:methanol:0.1 M ammonium acetate 100:100:4 (v/v/v) at a flow rate of 300 µL/min for 10 min. Levels of short-, medium-, and long-chain TAGs were calculated by referring to spiked internal standards of TAG (14:0)3-d5,TAG (16:0)3-d5 and TAG (18:0)3-d5 obtained from CDN isotopes, respectively. DAGs were quantified using d_5_-DAG17:0/17:0 and d_5_- DAG18:1/18:1 as internal standards (Avanti Polar Lipids).

Free cholesterol and cholesteryl esters were analyzed under atmospheric pressure chemical ionization (APCI) mode on a Jasper HPLC coupled to Sciex 4500 MD as described previously, using d_6_-cholesterol and d_6_-C18:0 cholesteryl ester (CE) (CDN isotopes) as internal standards^67^.

The lipidomic analysis employed a high-coverage, precise relative quantification workflow, the reliability and reproducibility of which have been well validated in previous studies^64–66^. Raw lipidomic data were first normalized using molar fractions to polar lipids (MFP). The polar lipid categories used for MFP normalization were PG, BMP, SL, CL, PE O, PE, PI, GM3, PS, LPI, LPE, LPA, PA, LPS, LPG, LPC, PC O, PC, SM, Sph, Cer, GlcCer, GalCer, and Gb3.

For multivariate analysis, principal component analysis (PCA) was performed using the *FactoMineR* package, and PCA plots with 95% confidence ellipses were generated using the *factoextra* package.

All heatmaps were created with the *ComplexHeatmap* package following global Z-score standardization of lipid abundance profiles rather than row-wise normalization. This approach allows the relative abundance of each lipid subclass within a given group to be intuitively assessed by the color intensity across columns.

For statistical comparisons among multiple groups, the non-parametric Kruskal-Wallis test followed by Dunn’s post-hoc pairwise comparisons was conducted using the *rstatix* package. The top 10 lipids with the smallest *P* values from Dunn’s tests were selected for differential lipid forest plots, whereas the top four lipids with the most significant differences from Kruskal–Wallis tests were used for boxplots and bar charts. All statistical plots were generated using the *ggplot2* package.

Lipid diversity across differentiation stages was assessed using the Shannon-Wiener diversity index, calculated from normalized molar fraction data of all detected lipid species. Calculations were performed using the *vegan* package in R, and temporal changes in lipid diversity were compared across five developmental stages.

For temporal pattern classification, normalized lipid abundance profiles were subjected to soft clustering using the *Mfuzz R* package. Lipids were categorized into four dynamic clusters: monotonically decreasing, increase-then-decrease, decrease-then-increase, and monotonically increasing patterns across the differentiation time course.

All annotated lipid species were further stratified by total acyl chain length and degree of unsaturation into saturated (no double bonds), mono- and di-unsaturated (1–2 double bonds), and polyunsaturated (> 2 double bonds) groups. Stage-dependent shifts in chain length distribution and unsaturation levels were statistically evaluated across differentiation stages.

Differential lipid species between pairwise differentiation stages were identified using volcano plot analysis. Significance thresholds were defined as |log₂(fold change)| > 1.0 and Benjamini-Hochberg false discovery rate (BH-FDR)-adjusted *p* < 0.01. Statistical testing was performed using two-tailed unpaired Student’s *t*-test followed by BH-FDR correction for multiple testing.

For volcano plot comparisons of lipid species, a more stringent significance threshold of BH-FDR-adjusted *p* < 0.01 combined with |log₂(fold change)| > 1.0 was applied to minimize false positive discoveries. This approach improves the clarity of the volcano plots by highlighting the most biologically meaningful lipid alterations and reducing visual clutter from marginally significant changes. Full lipidomic quantification for all detected lipid species are provided in Supplementary Table S3.

### Gene Ontology (GO) enrichment analysis

To minimize false positive enrichment results caused by unexpressed genes, we avoided using the whole genome as the background. Instead, strictly following established best practices for RNA-seq-based enrichment analysis^68,69^, we defined a custom background gene set consisting only of protein-coding genes that met the detection threshold (mean transcript per million [TPM] ≥ 1 in at least one developmental stage). Over-representation analysis (ORA) for GO terms was then performed using this customized background to ensure the statistical rigor of the enrichment calculations. We applied the Benjamini-Hochberg (BH) procedure to control the false discovery rate (FDR). For visualization and rigorous interpretation, enrichment plots were generated strictly based on FDR (BH-adjusted p-values) to illustrate the gradient of enrichment. Only terms with FDR < 0.05 were considered statistically significant.

### Quantification and statistical analysis

All experiments were performed with no fewer than three independent biological replicates. Quantitative data are presented as mean ± SEM, with n representing independent biological replicates. All statistical analyses were conducted using R (version 4.2.0) and GraphPad Prism (version 6). The specific statistical test used for each figure panel is explicitly stated in the corresponding Figure Legends.

Prior to statistical testing, the assumptions of normality and homogeneity of variance were assessed using the Shapiro-Wilk test. Statistical methodologies were applied based on data distribution and dimensionality.

### High-dimensional omics and functional enrichment

For high-dimensional pairwise comparisons of lipid species, the Benjamini-Hochberg (BH) method was applied to control the false discovery rate (FDR). For transcriptomic and epigenomic functional enrichment analyses (GO term analysis), p-values were adjusted using the BH-FDR method, with statistical significance defined strictly as FDR < 0.05.

### Multi-group comparisons

For normally distributed data involving more than two groups, a one-way ANOVA test was applied. For non-normally distributed multi-group data, the non-parametric Kruskal-Wallis test followed by Dunn’s post-hoc test with Bonferroni adjustment was utilized.

### Targeted and single-parameter comparisons

For targeted multi-gene qPCR panels (pre-selected based on prior omics screening) and other normally distributed single-parameter pairwise comparisons, two-tailed unpaired Student’s *t*-tests were performed.

Statistical significance was defined as *p* < 0.05 (or FDR < 0.05 where applicable). Significance levels are denoted as: ns, *p* > 0.05; * *p* < 0.05; ** *p* < 0.01; *** *p* < 0.001.

## SUPPLEMENTAL INFORMATION

Document S1. includes Figures S1–S4, related to Figures 1–7 and STAR Methods. Supplementary Table S1. Gene list of Cluster1.GOBP, related to Figure 6G.

Supplementary Table S2. Expression of cardiomyocyte differentiation related genes based on RNA seq data, related to Figure S1A.

Supplementary Table S3. Lipid analysis dataset, related to Figure 1E. Supplementary Table S4. Source data for heatmap, related to Figure 6H.

Supplementary Table S5. Integrated ATAC-seq and RNA-seq dataset, related to Figure 7B. Supplementary Table S6. Expression profiles of lipid-related genes, related to Figure S4B-F. Video S1. MLC2v-EGFP signals in beating Day-30 CMs, related to Figure S1.

## Notes

### Competing Interest Statement

The authors have declared no competing interest.

### Summary of Updates

In this revised version, we have mainly updated the Supplementary Results. In Fig. S1, we provide additional evidence supporting the maturity of day-30 cardiomyocytes, including immunofluorescence staining and electrophysiological assessment (Fig. S1E, F). The manuscript has also been thoroughly updated. In addition, the paper has been accepted by iScience (Tang Haite et al., iScience, 2026).

