## Supplementary Figures for "Lipid Metabolism Remodeling in Human Cardiomyocyte Differentiation and Maturation"

#### Document S1 includes Supplementary Figures 1 – 4

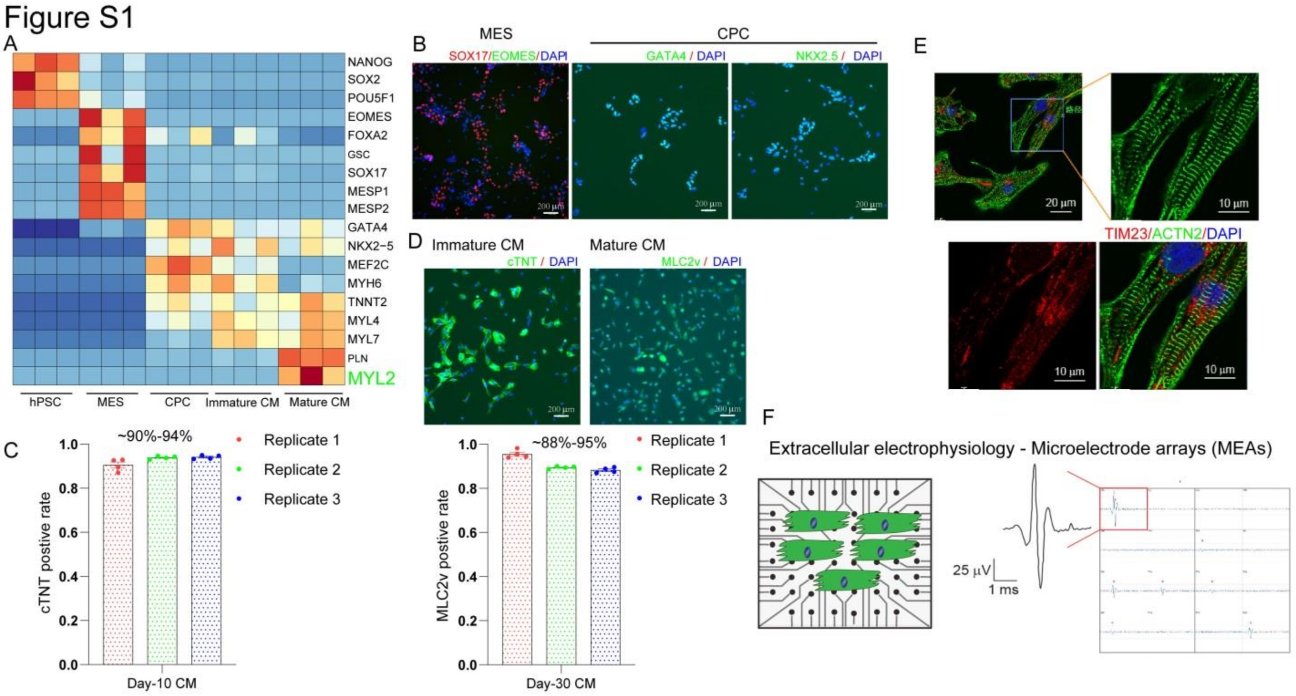


##### Figure S1. Validation of hESC-Derived CM differentiation Stages

(A) Quantitative bar plots validating stage-specific expression patterns of canonical pluripotent, mesodermal, cardiac progenitor, and cardiomyocyte developmental marker genes across five differentiation stages, based on RNA-seq data. *n* = 3 independent biological replicates per time point; data are mean ± SEM. Statistical significance was determined by one-way ANOVA followed by Tukey's post-hoc test.

(B) Representative immunofluorescence images illustrating sequential morphological and phenotypic changes during hESC-derived CM differentiation. Scale bar, 200 µm.

(C) Quantification of TNNT2-positive cell percentage in Day-10 CMs, counted from low-magnification immunofluorescence fields. Data are presented as mean ± SEM; each dot represents one independent biological replicate (*n* = 3), with 4 randomly selected fields analyzed per replicate.

(D) Quantification of MLC2v-positive cell percentage in Day-30 CMs, counted from low-magnification immunofluorescence fields. Scale bar, 200 µm. Data are presented as mean ± SEM; each dot represents one independent biological replicate (*n* = 3), with 4 randomly selected fields analyzed per replicate.

(E) Representative confocal immunofluorescence images showing sarcomeric organization in Day-30 CMs. Cells were co-stained for TIM23 (red, a vital protein component of the mitochondrial inner membrane) and ACTN2 (green, sarcomeric Z-line marker). Magnified views demonstrate highly ordered periodic sarcomeric striations. Scale bars: 20 µm (overview); 10 µm (magnified panels).

(F) Electrophysiological recordings of intracellular activity in Day-30 CMs were performed using the MaxOne single-well high-density microelectrode array (Maxwell Biosystems). Data are presented from independent biological replicates (*n* = 3).

### Figure S2

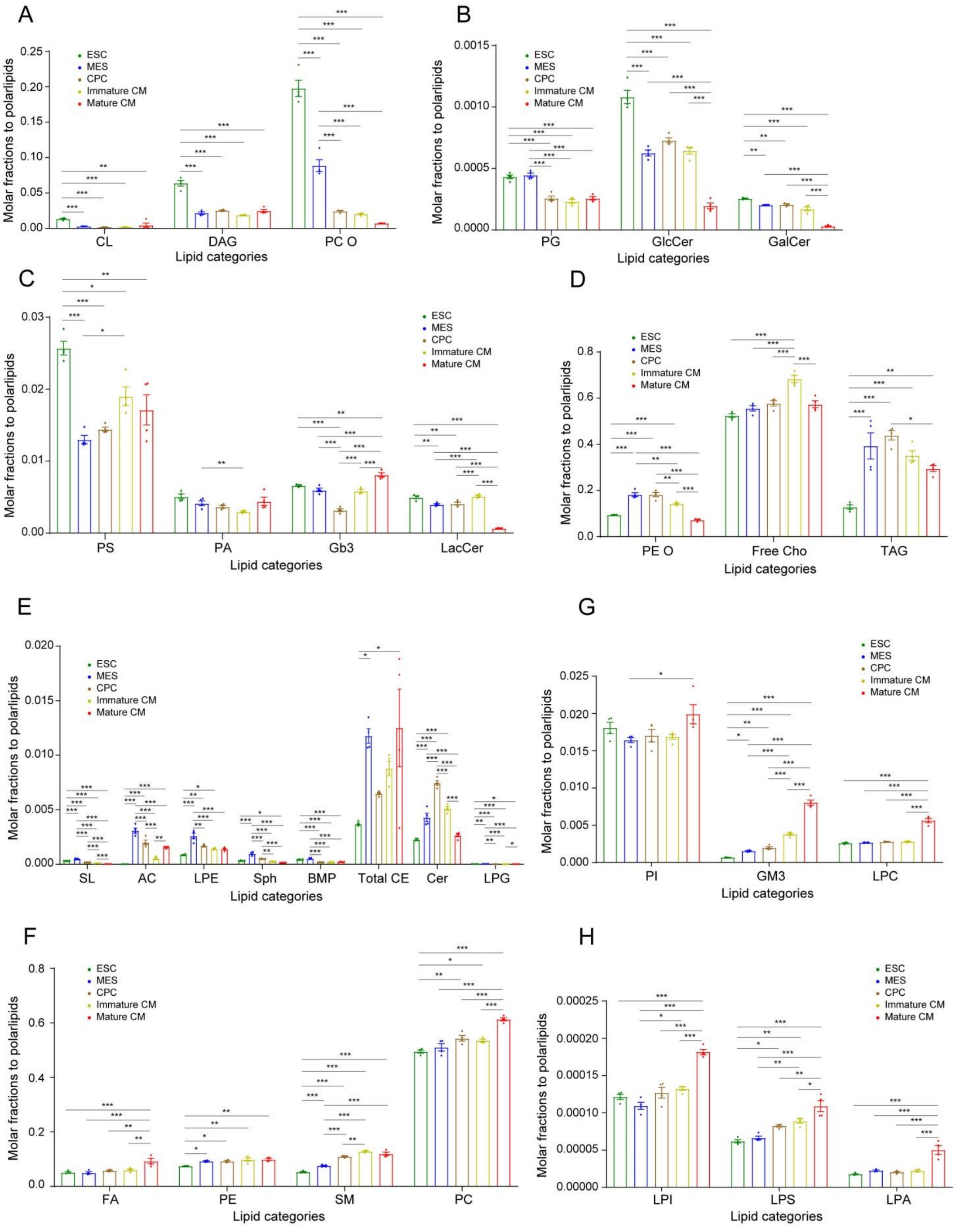


**Figure S2. Relative Abundance Dynamics of Major Lipid Classes during hESC-Derived CM differentiation**

(A)–(H) Quantitative bar graph validation of relative molar fractions for all 31 annotated major lipid categories across five developmental stages, corresponding to the global lipidomic heatmap in Figure 1E. Data are presented from independent biological replicates (*n* = 3). Statistical significance was determined by one-way ANOVA followed by Tukey's post-hoc test.

### Figure S3

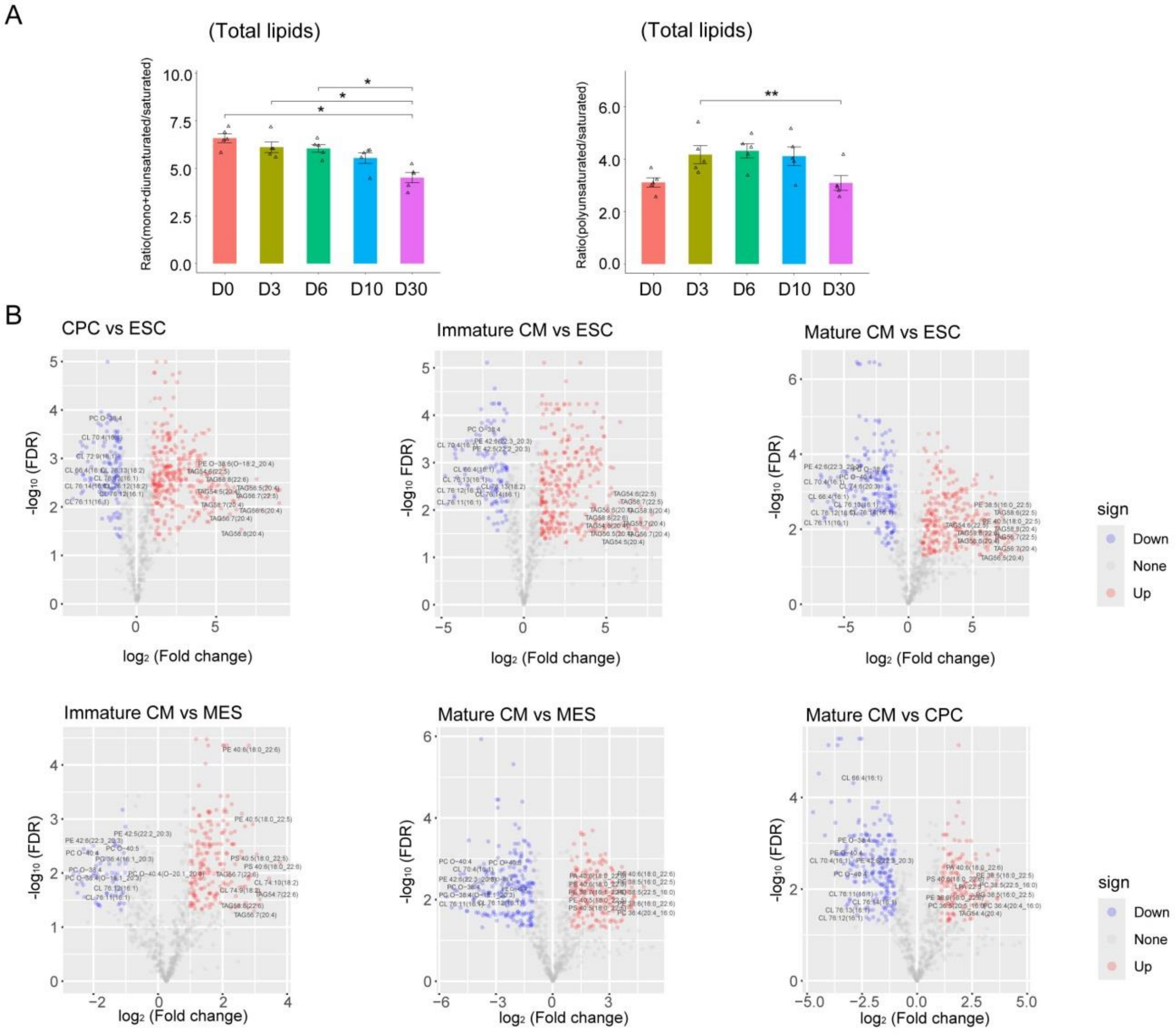


**Figure S3. Abundance Dynamics of Major Lipid Classes and Lipid Unsaturation Remodeling during hESC-Derived CM differentiation**

(A) Stagewise molar ratio comparison of unsaturated (mono+di, polyunsaturated) lipids relative to total saturated lipid species throughout differentiation. Data are presented from independent biological replicates ($n$ = 5); data are presented as mean ± SEM. Statistical comparison was performed using one-way ANOVA followed by Tukey's post-hoc test.

(B) Supplemental volcano plots showing pairwise lipid abundance comparisons across all non-consecutive developmental stages, including ESC vs CPC, ESC vs immature CMs, ESC vs mature CMs, MES vs immature CMs, MES vs mature CMs, and CPC vs mature CMs. These panels serve as a full data complement to the consecutive-stage comparisons presented in Figure 6E. Red dots denote significantly upregulated lipid species; blue dots denote significantly downregulated species. Significance cutoff: $|\log_2(\text{fold change})| > 1.0$ and BH-FDR adjusted $p < 0.05$, assessed via two-tailed unpaired Student's $t$-test followed by BH-FDR correction for multiple testing. Data are presented from independent biological replicates ($n >= 3$).

### Figure S4

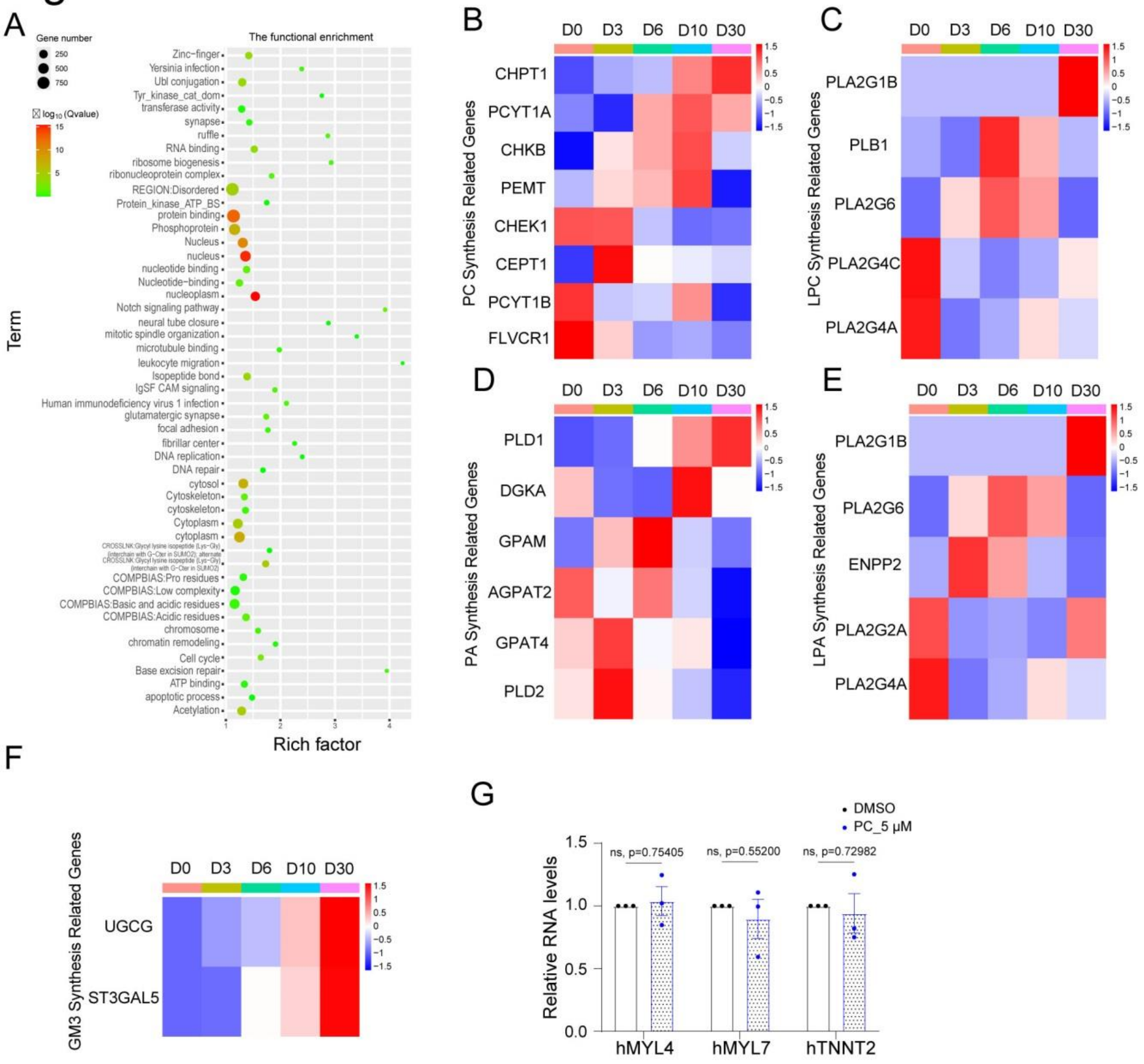


**Figure S4. Expression Dynamics of Lipid Metabolic Genes during hESC-Derived CM Differentiation**

(A) Integrative multi-omics correlation analysis of dynamic chromatin accessibility changes to temporal gene expression profiles of core lipid metabolic regulators during CM differentiation.

(B)–(F) Temporal expression trajectory heatmaps of key enzyme genes responsible for PC, LPC, PA, LPA, and GM3 lipid biosynthesis across all developmental stages.

(G) Transcriptional expression comparison of early cardiac lineage specification markers in immature CMs under 5 μM PC supplementation versus DMSO control. Data are presented as mean ± SEM from independent biological replicates ($n$ = 3). Statistical difference between two groups was evaluated using two-tailed unpaired Student's $t$-test.
